# Predicting cell morphological responses to perturbations using generative modeling

**DOI:** 10.1101/2023.07.17.549216

**Authors:** Alessandro Palma, Fabian J. Theis, Mohammad Lotfollahi

**Affiliations:** Institute of Computational Biology, Helmholtz Center Munich – German Research Center for Environmental Health, Neuherberg, Germany; Wellcome Sanger Institute, Wellcome Genome Campus, Cambridge, UK; Department of Mathematics, Technical University of Munich, Munich, Germany; School of Life Sciences Weihenstephan, Technical University of Munich, Munich, Germany

**Author notes:** co-senior authors (Fabian J. Theis, Mohammad Lotfollahi).

## Abstract

Advancements in high-throughput screening have enabled the exploration of rich phenotypic readouts like high-content microscopy, expediting drug target identification and mode of action studies. However, scaling these experiments to the vast space of drug or genetic manipulations poses challenges, as only a small subset of compounds show activity in screenings. Despite being widely used in various applications, machine learning methods have not shown a reliable ability to extrapolate predictions to scenarios involving unseen phenomena, specifically transforming an unseen control cell image into a desired perturbation. We present a generative model, the IMage Perturbation Autoencoder (IMPA), which predicts cellular morphological effects of chemical and genetic perturbations using untreated cells as input. IMPA learns perturbation-specific styles from generalized embeddings and generates counterfactual treatment response predictions in control cells. We demonstrate IMPA can predict morphological changes caused by small molecule perturbations on breast cancer cells. Additionally, we test IMPA on the unseen drug effect prediction task, showing improved performance over state-of-the-art generative models when compounds are structurally related to the training set. Finally, generalizability and capability to predict more subtle effects are showcased through its application to large microscopy datasets with hundreds of genetic perturbations on U2OS cells. We envision IMPA to become a valuable tool in computational microscopy for aiding phenotypic drug discovery, facilitating navigation of the perturbation space, and rational experimental design.

## Introduction

The advent of high-throughput image-based profiling has made it possible to screen thousands of chemical [1–4], and genetic perturbations [5, 6] to identify potential drug targets, modes of action (MoA) and gene functions. Such profiling assays measure morphological changes of sub-cellular structures by staining them using multiple fluorescent dyes [7] to highlight different organelles and components. An example is Cell Painting [8], the most commonly used unbiased image-based profiling assay [9] to measure different substructures using fluorescent dyes. While current technologies are scalable, it is unfeasible to explore the combinatorially large space of potentially synthesizable drug molecules or genetic perturbations [10, 11]. Consequently, performing larger screens can be experimentally challenging and costly. To improve navigation of the perturbation space for an optimal experimental design, computational methods are needed to predict morphological responses to perturbations not measured in the experiment. The generated cellular responses can be used to narrow down the hypothesis space facilitating phenotype-based drug discovery [12].

Computational approaches for predicting phenotype responses in high-throughput image-based data have been explored in supervised and unsupervised settings. Supervised tasks include compound mode of action (MoA) [13–18] and drug toxicity [19] prediction, and assay activity annotation [20]. In contrast, unsupervised methods can be used to generate in-silico representations of cell image features under specific interventions and conditions [21] or predict (polypharmacological) responses to drug combinations [22]. However, the field currently lacks a generative model that provides counterfactual predictions on perturbation images. In the context of generative modeling for biological data, MorphNet, a Generative Adversarial Network (GAN) model, synthesizes cell morphology based on gene expression but does not tackle perturbation-specific image generation [23]. Other recent studies focus on representation learning. For example, CLOOME [24] learns a joint representation of images and drug molecular structures via contrastive learning and exploits the embedding space for drug- or image-conditioned instance retrieval. However, the model is not designed to predict perturbation responses in control cells. Moreover, concurrently with our work, generative modeling for domain adaptation has been utilized to create batch-invariant representations of cells in high-content microscopy screenings [25], yet its application to perturbation prediction on cell morphology remains unexplored. Meanwhile, Mol2Image employs a conditional flow-based generative model to generate cell images based on chemical compound structures. However, it lacks a detailed exploration of the synthesized images of unseen drugs and does not account for perturbation-induced morphological changes in control cells [26].

In summary, this gap prompts a new approach to answer the question: “How would an image of an untreated cell appear if it had been subjected to a specific perturbation?”. To address this need, we introduce the Image Perturbation Autoencoder (IMPA), a deep generative model designed to predict cellular responses to small molecules and genetic perturbations in high-throughput image-based profiling screens. IMPA adopts a style transfer approach [27–30] for the image-to-image translation [31, 32] task. The model learns to decompose a cell image into its style (perturbation representation) and content (cell representation). Through training, IMPA can transfer a cell to a desired style i.e., perturbation, while preserving its perturbation-independent content. IMPA stands out for its utilization of unpaired data, removing the requirement to screen images of the same cell before and after treatment. Considering the impracticality of collecting paired images in large-scale experiments, this aspect is crucial. We demonstrate the effectiveness of IMPA on diverse datasets, including drug screens on MCF-7 cells and hundreds of RNA interference (RNAi) perturbations on U2OS cells. Furthermore, we showcase IMPA’s interpretability and flexibility by interpolating across styles and contents, observing the intermediate perturbation effects, and generalizing to compounds not seen during the training. Our model enables efficient target discovery in phenotypic screens, streamlining experimental design and enhancing our understanding of morphological alterations in high-throughput image-based profiling.

## Results

### Learning morphological responses to perturbations using style and content formulation

We model the phenotypic responses to perturbations in high-content imaging screens by decomposing the representation of each image into the perturbation it was treated with (i.e., *style*) and a representation of the cell (i.e., *content*). This approach is based on “style-transfer” [34], an active area of research in deep learning and computer vision [35, 36]. Style transfer involves modifying the characteristics (e.g., the art style) of an image to another while preserving the original content of the image. Following this approach, we developed IMPA, a conditional generative adversarial network [37] that, given an input image of control cells, can generate the counterfactual image response to a desired perturbation (see **Fig. 1a** or **Supplementary Fig. 1a-b**). IMPA is built upon the architecture proposed in StarGANv2 [38]. However, we applied modifications to the conditioning mechanism in a way that supports the usage of prior perturbation embeddings rather than Gaussian random style encodings like in the original model (see **Architecture**). This modeling aspect provides flexibility in the choice and representation of perturbations for IMPA, allowing for diverse examples such as incorporating molecular structures for drug screening and co-expression-based gene representations for gene knockdown studies as styles for perturbation modeling.

**Figure 1.**
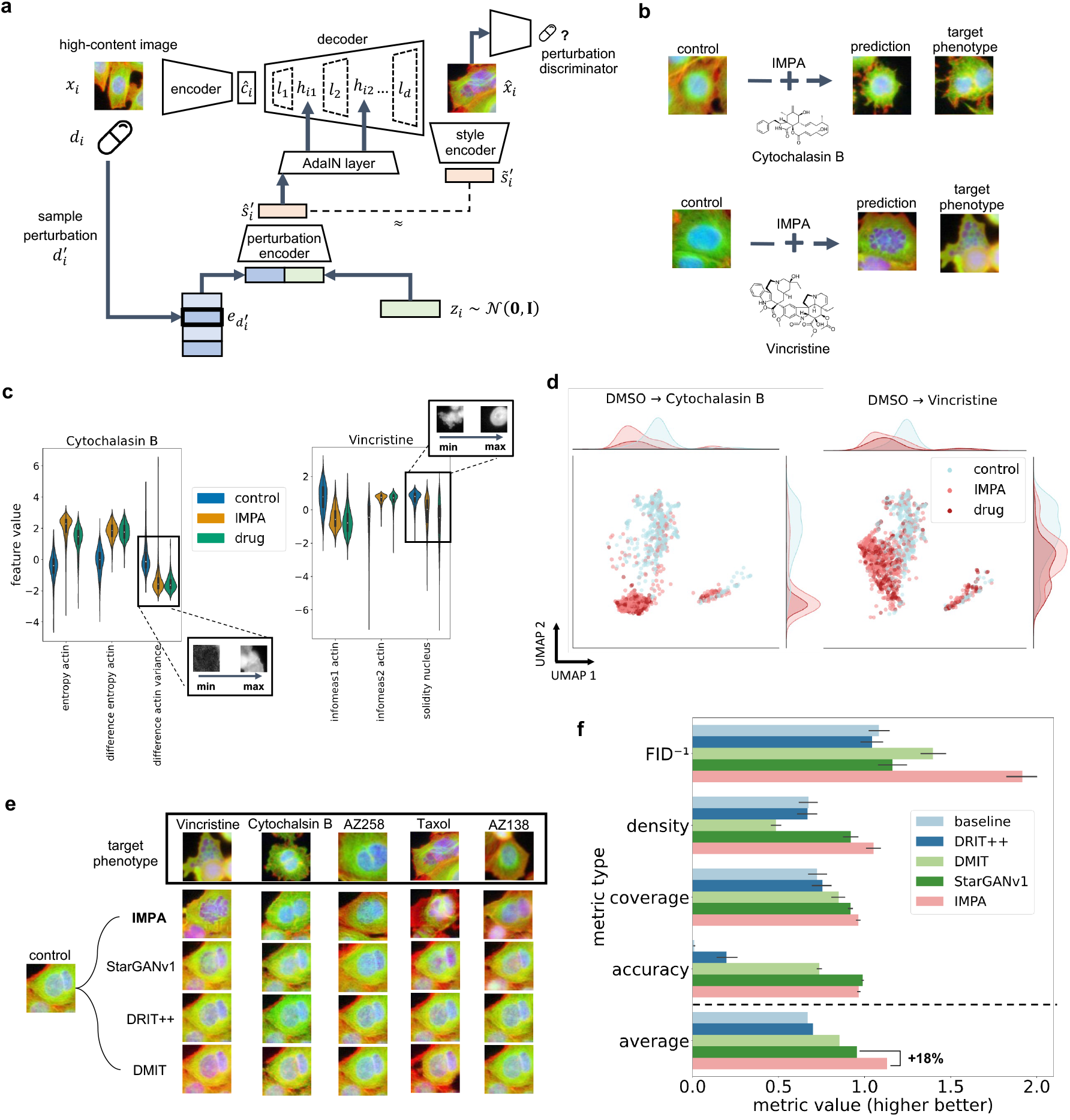
IMPA enables perturbation effect prediction via style transfer. **(a)** A visual description of the architecture of IMPA. A control cell image x*_i_* is encoded into a content representation. When applying transformations to the input, a dense embedding of the target perturbation is collected and concatenated with a normally-distributed random vector. A low-dimensional projection of the concatenation constitutes the style space which is used to condition every layer l*_i_* of the decoder via the AdaIN method [33]. With h*_ij_* we indicate the output of the j*^th^* decoder layer on image i. The transformed output is used to induce a discriminator net into predicting that the decoded image comes from the target perturbation batch. Moreover, a style encoder is trained to replicate the style vector from the transformed image. **(b)** Synthetic perturbation of control cells (left) with Vincristine and Cytochalasin B for a reduced version of the BBBC021 dataset (5 perturbations, 20,313 cell images). The generated images approximate the style (phenotype) of the ground-truth examples on the right. **(c)** Distribution of most important discriminative CellProfiler features (see the text) between controls, IMPA’s predictions, and original perturbation images for Vincristine and Cytochalasin B. Feature importance values were estimated with a Random Forest classifier trained to discriminate controls from perturbation images based on CellProfiler morphological measurements. **(d)** UMAP plots showing the distribution of CellProfiler morphological features as in (c) before and after transformation with IMPA for Vincristine and Cytochalasin B. **(e)** Visual Comparison of IMPA with existing models on the style transfer task. The models are used to transform a control cell image (left) into its perturbed counterparts (right) for each drug. **(f)** Evaluation metrics comparing generated images with the real perturbed images. Results are averaged across drugs and 10 bootstrap repetitions. For each measurement, the higher the value, the better the model approximates real perturbation images. The FID score was divided by 100 and inverted to yield a range comparable with other metrics.

Given a dataset *{*x*_i_}^N^* of high-content images of cells and the associated perturbation index *{*d*_i_}^N^*, IMPA decomposes the latent representation of a cell x*_i_* into a content ĉ*_i_* and a style ŝ*_i_* component. The content ĉ*_i_* is inferred through a convolutional encoder that projects the input image onto a content space. To learn a representation for the style, we devised a perturbation and a style encoder (see **Fig. 1a** or **Supplementary Fig. 1a-b**) interacting together to learn an embedding for each perturbation. The perturbation encoder receives a concatenated vector of noise z*_i_* and an embedding e*_d_′* representing a perturbation d*^′^* different from the original d*_i_* and linearly projects said concatenation onto a lower-dimensional style ŝ*_i_*. The perturbation embedding choice is flexible. In our study, we employed physio-chemical descriptors [39] for representing drugs and Gene2Vec embeddings [40] for genetic perturbations as the inputs of the perturbation encoder. However, it is relevant to note that alternative options are available for the perturbation embeddings such as representations extracted from pre-trained models [41], information from knowledge bases [42, 43] or fully-trainable embeddings [44]. The probabilistic nature of the perturbation style allows us to learn a full range of possible responses to each treatment in the training dataset.

Adaptive Instance Normalization (AdaIN) [33] manipulates the content encoding to match the input perturbation by scaling and shifting feature maps in the convolutional decoder based on the conditioning style. During training, IMPA combines random style embeddings and the contents using AdaIN in each layer of the decoder (see **Fig.1a** and **Architecture** for details). In contrast with conditional generative approaches for perturbation prediction on single-cell RNAseq [41, 44], we opted for stepwise conditioning of the decoding process rather than altering the output of the encoder. This choice was made to accommodate the more complex nature of image generation.

The generated image is fed to a convolutional style encoder, which outputs a prediction s^∼1^*_i_* and is trained to approximate the output of the perturbation encoder ŝ*^′^*. Effectively, the style encoder links image features to a perturbation-specific style and can be used to inspect single-cell representations upon training. Such an approach enforces alignment between features extracted from the generated images and the learned style embedding, making them consistent. To ensure that predictions correctly match the desired perturbation style, we train a discriminator classifier [45] for each perturbation to discriminate between real images of cells treated with such a perturbation versus generated images. The decoder network has to synthesize accurate images of the effect of target d*^′^* to deceive a multi-task discriminator into classifying it as a true image from the target style (perturbation). Conversely, the discriminator is trained on real data to recognize the difference between a true and a generated treatment image. The multitask discriminator does not try to classify images between different perturbations but to predict whether an image is true or generated according to a perturbation class. This approach makes it more suitable when the phenotypic responses to different perturbations are similar and, as such, hard to classify. Finally, a cycle-consistency loss [46] is employed to encourage the model to learn reversible mappings (see **Supplementary Fig. 1b**), hence, transferring perturbed images back to the style of the input condition. The interplay of all units was shown to significantly improve generative performance in StarGANv2 [38].

### IMPA accurately predicts morphological changes after drug perturbations

We evaluated IMPA on predicting morphological changes after drug perturbation using the BBBC021 dataset (see **Datasets**) [47]. BBBC021 comprises images of p53-wildtype breastcancer model cells (MCF-7) perturbed with 113 compounds and imaged across three channels: nucleus, β-tubulin and actin. We employed a widely-used reduced version of the dataset from [13] comprising 38 compounds with expected phenotypic effects at the tested concentrations. We pre-processed the dataset from whole slides to images cropped around single cells, retrieving a total of around 97k image patches. For a more straightforward assessment of generated images, we first trained our model on a subset of five drugs (AZ138, AZ258, Cytochalasin B, Taxol, Vincristine) with visible effects on the morphology compared to controls according to [47] (see **Supplementary Fig. 2a**). For this scenario, a perturbation embedding for each drug was extracted using the RDKit software package [39]. Each embedding represents descriptors that define topological and structural drug properties derived from atomic neighborhoods in the molecular graph [48].

Qualitative analysis of predictions showed that IMPA transformed control cells to target perturbation styles while preserving content information such as cell orientation and translation (see **Fig. 1b** and **Supplementary Fig. 3c**). For example, it is known that Cytochalasin B blocks actin polymerization [49]. IMPA’s prediction for Cytochalasin B resulted in the loss of homogeneity and extension in the actin channels compared to the source control image. Moreover, the predictive capabilities of IMPA were demonstrated by its ability to accurately anticipate the expected loss of nuclear integrity caused by Vincristine [50] (see **Supplementary Fig. 3a** for more examples).

To assess the impact of IMPA on important features characterizing perturbations, we employed the CellProfiler software [51] to compute a series of 356 features, including shape, texture, area, and intensity distribution of cell images before and after transformation by IMPA. Our objective was to investigate how IMPA affects these features and their alignment with actual perturbed images compared to control cells (see **Fig. 1c** and **Supplementary Fig. 5c**). To subset relevant features for each perturbation, we estimated feature importance by training a Random Forest classifier [52] using CellProfiler features to discriminate control cells from perturbed cells in separate binary classification settings. The distribution of important morphological features in transformed images showed a similar shift to actual images when compared to controls (shown as the baseline in **Fig. 1c**). For example, we can see an increase in actin entropy in Cytochalasin B (actin disruptor) or a loss of nuclear solidity in Vincristine (tubulin destabilizer). Additionally, dimensionality reduction via UMAP on the features extracted from real and transformed images revealed that CellProfiler features of generated images aligned better with real perturbed images than controls, corroborating our observation that the model alters biological features coherently with the expected perturbation outcome (see **Fig. 1d** and **Supplementary Fig. 5b** for all five drugs).

We further compared IMPA with three existing GAN models performing the style transfer task (see **Baselines**). Notably, these prior models were not specifically designed or utilized in the context of conditional generation with prior perturbation embeddings. Unlike IMPA, which incorporates a multi-task discriminator to guide the generation process, the other models adopt a multi-class convolutional classifier as the conditioning mechanism. Additionally, one of the compared models, StarGANv1 [53], is deterministic and, therefore, does not learn a distribution of responses to drugs. We hypothesized that these design choices may limit the performance of these models, particularly when perturbation effects are subtle or noisy. Qualitatively, our visual analysis revealed that DRIT++ [54] did not exhibit significant deviation from the source images, while StarGANv1 and DMIT [55] successfully altered intensity distributions but tended to over-preserve control morphology (see **Fig. 1e**).

To quantitatively compare these models, we employed three commonly-used [56] GAN evaluation metrics: Fréchet Inception Distance (FID) [57], Density, and Coverage [58], which assess the similarity between the distribution of generated images and the target real image distribution. In addition, we measured an accuracy metric that evaluates how often a pre-trained classifier correctly labels generated images with the mode of action known for the target drug (see **Evaluation metrics** for a detailed description of the evaluation process). To establish a lower-bound performance, we included a baseline comparison between control cells and real perturbed images, simulating a model incapable of style transfer. Our findings demonstrated that IMPA outperformed the compared style transfer models in three out of four metrics (see **Fig. 1f**). Specifically, IMPA achieved a 27% decrease in FID (showed as its inverse in **Fig. 1f**), a 5% increase in coverage, a 14% increase in density, and an overall 18% performance improvement compared to the second-best method for each metric. Regarding mode of action classification accuracy, IMPA performed comparably well to StarGANv1, exhibiting only a 2% decrease in classification accuracy, while significantly outperforming all other models. It is worth mentioning that we did not conduct a direct comparison between IMPA and StarGANv2. This decision was based on the similarity between the discriminator-generator structures of StarGANv2 and our model, which makes a direct comparison less informative in this context.

### IMPA enables extrapolation to unseen drugs

A long-standing challenge for drug screening is selecting a chemical library to cover different areas of the chemical space [10, 59, 60] corresponding to diverse modes of action. However, fully exploring all possible compounds is impractical due to experimental costs and logistics [61, 62]. Therefore, in-silico methods are needed to predict the response to unmeasured compounds [41, 44]. IMPA’s architecture allows for tackling this challenge by simply using chemical representations for unseen drugs at test time.

Interpolation between drug effects and extrapolation to unseen treatments are possible only if the model learns a smooth style space reflecting the phenotypic heterogeneity present in the dataset. A meaningful style encoding can be used to navigate the chemical space to gain insight into drugs with similar effects or intermediate phenotypes between untreated and treated cells (see **Fig.2a** and **Supplementary Fig.8**). To show that the style space is amenable to interpolation, we studied the effect of generating images of cells from drug encodings lying in the linear path between the embeddings of controls and target drugs. Style space interpolation may offer insights into various aspects such as response similarity to multiple perturbations or differences in the effect size of drugs with similar modes of action on the phenotypic spectrum. However, further research is needed to fully explore and understand the implications of drug space traversals and their relationship with morphological changes. We inspected the results across the trajectory by classifying intermediate images into training drugs with a classifier trained to discriminate between perturbation images. For drugs such as AZ258 (see **Fig. 2a**), the classifier correctly assigned untransformed images to controls and fully-transformed images to the target drug class. The experiment suggested that, before yielding the final target phenotype, the classifier is uncertain between labeling transformed images as AZ258 or Mitomycin C, while adding the whole drug style generates reliable AZ258 images. When training on large screenings, IMPA can gradually perturb control images with different levels of a drug style and explore the chemical space around unseen treatments to define potential similarities with molecules of known impact on cell morphology.

**Figure 2.**
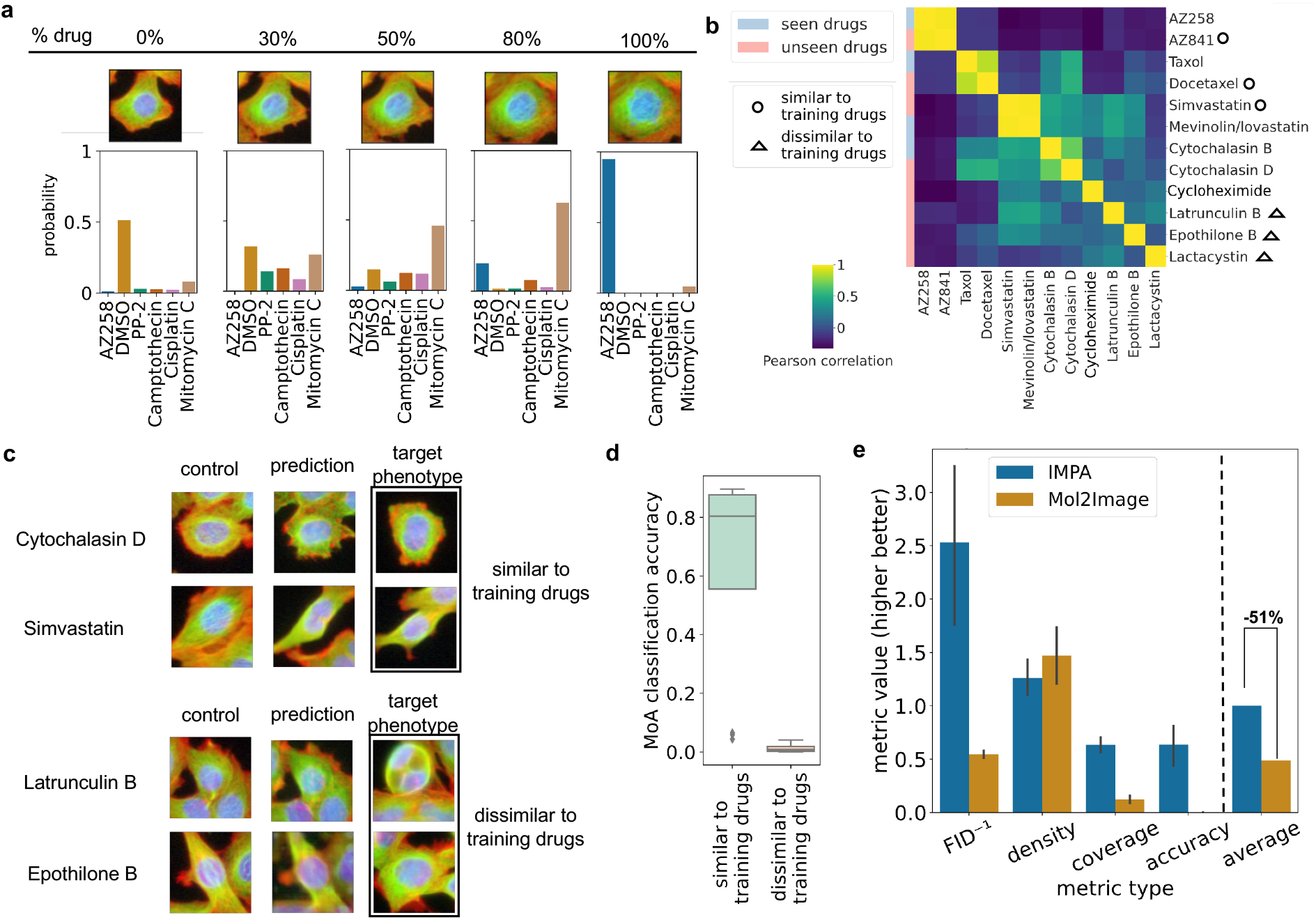
IMPA predicts perturbation responses to unseen drugs. **(a)** Examples of style interpolation between controls and the training drug AZ258. A classifier pre-trained to recognize training perturbation images is applied to each interpolation step to investigate the similarity to existing compounds when adding different levels of drug style to the DMSO. **(b)** Heatmap displaying the correlation between the RDKit embeddings of pairs of drugs from a subset of BBBC021. The row color indicates if the drug is selected as an unseen example or included in the training process. **(c)** Generated images for unseen drugs are divided into two groups. Similarity to training drugs or lack thereof is assigned based on RDKit embedding correlation with other drugs with the same MoA in the training set. **(d)** Frequency with which the MoA of the target unseen perturbation is assigned to transformed images by a pre-trained classifier. Separate scores are derived for drugs similar and dissimilar to training compounds. The higher the score, the better the prediction. **(e)** Comparison between IMPA and Mol2Image on the unseen drug effect prediction task. The metrics are evaluated on four compounds: AZ841, Docetaxel, Simvastatin, and Cytochalasin D. For all measurements, the higher the value, the better the generated output approximates the expected phenotype.

We employed IMPA to predict responses to previously unseen drugs to investigate the potential of utilizing prior perturbation embeddings for conditioning style transfer. In this section, we extend our analysis to the entire BBBC021 dataset, consisting of a cohort of 34 drugs (derived by excluding 4 drugs with undisclosed chemical structures). Unlike the subset used in the previous section, this comprehensive dataset provides a broader representation of the chemical space. A subset of samples from eight distinct treatments was deliberately withheld from the training process. This set of eight drugs was carefully selected to encompass both a group of drugs structurally correlated or functionally associated with at least one training compound (AZ841, Docetaxel, Cytochalasin D, Mevinolin) and compounds whose feature representation is not correlated to any of their functionally-related drugs (Epothilone B, Latrunculin B, Cycloheximide, Lactacystin) (see **Fig. 2b**). For brevity, we refer to the former set as *similar to training drugs* and the latter as dissimilar to training drugs. We hypothesized that IMPA generalizes well only on the former category as it fits a chemical space where similar embeddings induce a similar phenotypic response. Comparing predictions for these two groups of drugs confirmed our hypothesis that IMPA can more accurately predict response to drugs with higher similarity to training compounds (see **Fig. 2c** and **Supplementary Fig. 7d**). For example, IMPA visually generated actin degradation traits from Cytochalasin D thanks to its similarity to Cytochalasin B. Conversely, the model produced generic outputs for drugs with structures dissimilar to those seen in training compounds and failed to approximate their target phenotypes. We corroborated our observation by applying a pre-trained MoA classifier to predictions for unseen drugs (see **Fig. 2d**). Our results showed that, while an average of 63% of generated samples from unseen drugs similar to the training treatments is correctly classified with its ground-truth MoA, IMPA manages to model the phenotypic response to drugs of the dissimilar group less than 2% of the time. These results demonstrated IMPA’s potential failure cases and limitations, essential to understanding when to apply in-silico models.

In this study, we directly compared our approach, IMPA, to Mol2Image [26], a flow-based generative model that generates images of perturbed cells conditioned on learned graph-based drug representations. The core difference is that Mol2Image generates images of perturbed cells conditionally from Gaussian noise, while IMPA performs style transfer on existing images conditioned on prior perturbation embeddings. Thus, our approach has the advantage of enabling the study of differential morphology between control and perturbed conditions, providing a deeper understanding of the impact of drug-induced changes on cellular phenotypes. We benchmarked both models by leaving out samples from the four compounds with higher similarity to training drugs, as reported in the previous section (see **Fig. 2e**). Overall, we observed that Mol2Image’s average performance across the evaluation metrics mentioned earlier amounted to approximately half that of IMPA when predicting the effect of unseen drugs similar to the training compounds. This gap was particularly evident in FID and coverage. Specifically, the latter metric estimates the portion of a target condition’s space recovered by the generated image samples. Poor performance of Mol2Image on such an evaluation criterion implies a lack of exploration of the whole response spectrum to a perturbation, which is, on the other hand, successfully tackled by IMPA.

### IMPA predicts phenotypic responses to genetic perturbations

To evaluate the generalization capabilities of IMPA and test its ability to tackle challenging settings with more subtle morphological effects, we conducted experiments on two gene-knockdown datasets using U2OS cells (see **Datasets**). The first dataset, BBBC025, comprised 350 distinct single-gene perturbations achieved through shRNA-mediated RNA interference targeting 41 different genes (see **Supplementary Fig. 9a**). The second dataset, RxRx1, consisted of 1,138 siRNA perturbations (see **Supplementary Fig. 10a**). All perturbations were tested in multiple experimental batches. In line with the drug experiments, we employed a dense representation of the genes targeted by the genetic perturbations as a proxy for the style transfer process. As an embedding for the target genes, we selected the Gene2Vec model [40], which provides a vector representation of genes based on co-expression patterns detected in large cohorts of expression datasets. This resulted in 41 different style representations for BBBC025 and 1,070 for RxRx1 after selecting the siRNAs whose target genes have known Gene2Vec embeddings. For control instances, we utilized a designated token with its corresponding embedding.

To assess the representation potential of our model, we generated an embedding for each perturbation by extracting image features using the style encoder and computing their average to obtain a single point per perturbation. These representations were then visualized in a two-dimensional space (see **Fig. 3a-b**). To assess whether the distribution of such perturbation representations is associated with effect size, we annotated the average distance between each perturbation and their respective controls on the two-dimensional UMAP coordinates. Since images of perturbations and controls are unpaired, we consider the FID between controls and perturbations to define the response size. Such a score is used to compute the perceptual difference between two sets of images as the Wasserstein distance of their feature representations according to a pretrained convolutional network [57, 63]. Our results exhibited a cohesive gradient on the UMAP plot, especially in RxRx1. More specifically, the average style encoding of impactful perturbations gradually separated from those resembling controls, which in turn clustered together (see **Fig. 3a-b**). Furthermore, the findings suggested that IMPA effectively predicted both datasets’ phenotypic effect of active knockdowns when we selected perturbations with the highest distance from the control. Conversely, other image-translation models such as StarGANv1 and DRIT++ failed at this task, either producing subpar results or not capturing the morphological deviations caused by active perturbations (see **Supplementary Fig. 9-10**).

**Figure 3.**
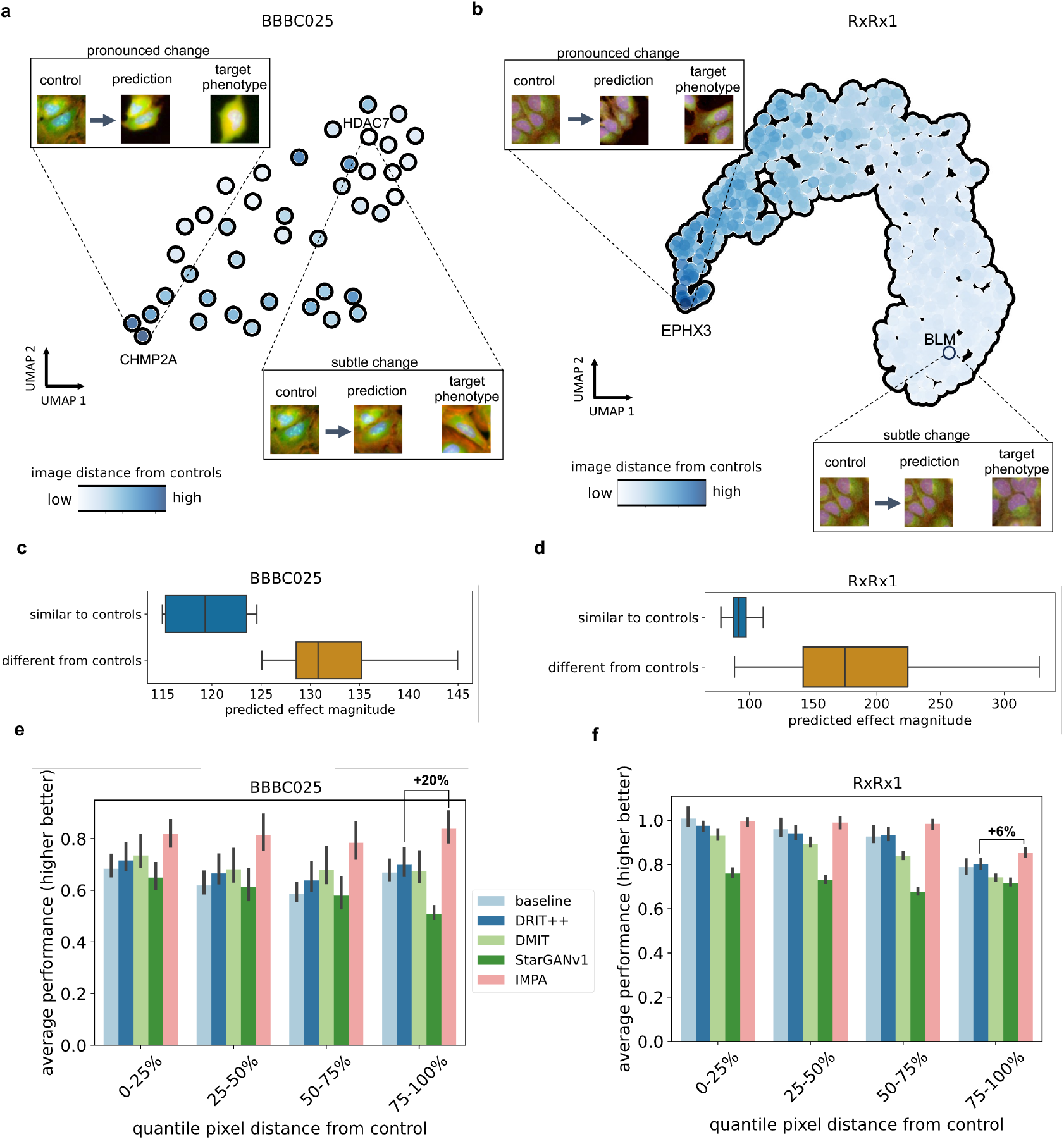
IMPA predicts morphological response to hundreds of genetic perturbations. **(a-b)** UMAP plot of the mean style embeddings extracted from perturbation images for BBBC025 (41 gene targets, 101,100 images) and RxRx1 (1,070 gene targets, 170,943 images) datasets. Each point represents a perturbation whose images are embedded and averaged using the style encoder. Each data point is colored by the mean difference between the associated perturbation images and the respective controls measured as FID. We additionally report examples of IMPA’s transformations on the perturbations with the highest and lowest distance from controls. **(c-d)** Boxplot showing the magnitude of phenotypic change induced by IMPA as the distance between images before and after transformation. Controls are transformed into the 10% closest and furthest perturbations to the respective untreated cells in terms of FID. **(e-f)** Average prediction metrics for perturbations with different pixel distances from control images. On the x-axis, perturbations are grouped in percentile ranges of the average distance from the controls measured as FID (a higher percentile means a higher difference between the perturbation images and the control). On the y-axis, the average performance of style transfer models measured in FID, coverage, and density compared to ground-truth images.

In large-scale, unbiased phenotypic screens, it is anticipated that only a small portion of perturbations will yield a significant effect. Therefore, it is important for a generative model to capture both effect and absence thereof in its output. To investigate this, we quantified the distance between control images before and after transformation by IMPA in terms of FID (see **Fig. 3c-d**). We considered the 10% closest and furthest perturbations from the controls, expecting to notice a significantly stronger morphological shift for the latter class. As expected, for perturbations whose images were close to their respective controls, IMPA induced 10% and 50% average weaker morphological changes than the 10% furthest knockdowns from their untreated counterparts in BBBC025 and RxRx1, respectively. This observation suggested that the phenotypic transitions induced by our model are correlated with the actual perturbation effect size observed in the dataset.

Furthermore, we assessed the model’s performance as a function of the distance between the distributions of perturbation and control images in both the RxRx1 and BBBC025 datasets (see **Fig. 3e-f**). In our evaluation, we compared IMPA to previously described GAN models and the baseline of untransformed control images. When transforming controls into perturbations with negligible or no effects, we expected our model to predict the absence of a signal. Thus, a minor or no change should be applied to source images. In such a scenario, we anticipated a score similar to the baseline, which compares untreated cells with perturbed cells in genetic interventions. Conversely, the baseline was expected to yield inferior results for highly active perturbations compared to predictive models. This is because these models should change the source images to capture drastic changes in the target perturbations. In both datasets, for perturbations close to controls (within the 0-25% FID percentile), most models performed similarly to the baseline as they learned not to apply substantial morphological alterations to untreated cells. For IMPA this was particularly evident in the RxRx1 dataset, where the model’s output was very close to the untransformed images when no effect prediction was expected. Moreover, IMPA demonstrated less performance deterioration than the competing models when handling perturbations with the highest distance from their controls (within the 75-100% FID percentile). This aspect was pronounced in the BBBC025 dataset, where IMPA outperformed the second-best model by an average of 20% of its score. On RxRx1, DRIT++ exhibited similar performance to IMPA, with only a 6% average performance decrease.

Overall, our results validate the model’s scalability to large and noisy perturbation screens with diverse degrees of effect sizes and the ability of IMPA to fit a meaningful representation space that generalizes across various perturbation effects.

## Discussion

Synthetic prediction of morphological responses to perturbation in high-content screenings is paramount to orienting experimental design towards promising treatments. To this end, we implemented IMPA, a conditional generative adversarial network that morphs images of untreated cells into what they would look like under the effect of a queried perturbation. IMPA derives treatment-specific latent encodings and uses them as a style to overlay the phenotypic effect due to perturbations onto control cell images. More specifically, we interpreted the problem of *synthetically perturbing cells* as disentangling content from style information on input images. While the former informs the model on perturbation-agnostic features such as cell orientation and relative positioning in the plate, the latter induces a target phenotype. Notably, this approach differs from the previously proposed pipelines for disentangling the perturbation signal from the basal cell state [41, 44]. In fact, we found the style transfer paradigm coupled with GAN training is more effective on the output quality, as it allows keeping a high-dimensional content representation while relying almost completely on a low-dimensional style encoding to model morphological shifts.

We demonstrated the use of IMPA on different modalities of perturbation data. In drug screenings, we derived the style encoding to condition image translation from dense molecular embeddings extractable from any compound with a known structure. In this framework, we showed that IMPA successfully approximates target perturbations qualitatively and quantitatively and outperforms three other image-to-image translation models in the task. Moreover, IMPA’s ability to use structural information from drugs to condition perturbation style transfer allows the model to predict the phenotypic effect of unseen treatments as long as their representation is correlated with that of training compounds with a similar effect. We also showed that IMPA can be generally applied to other perturbation modalities (e.g., gene knockdown studies).

While IMPA performs well in the proposed settings, training deep generative models on modern large-scale perturbation datasets with millions of images [64] remains a challenge. Aside from the noisy response, IMPA does not account for batch effects and is therefore vulnerable to confounding sources such as the acquisition venue and the experimental batch when not all perturbations are tested in all technical conditions. To address this limitation, future work may focus on integrating batch information by enriching the conditioning mechanism with batch labels, enabling correction for confounding effects. One possible strategy to explore is the combination of our model with the approach proposed by Pernice et al. [25], which leverages style transfer for batch correction in high-throughput screenings. Moreover, generalization to unseen perturbations heavily depends on the embeddings employed to learn the style. Finally, it should be stressed that IMPA’s prediction on unseen perturbations is successful only when the model has seen a similar compound with the same effect in the training compounds. This makes IMPA more suited to identify novel compounds with known effects rather than uncovering new phenotypes.

We envision future work investigating the relationship between the chemical space and morphological changes more deeply. This will involve exploring how different regions of the chemical space are associated with specific morphological transformations. By understanding this relationship more deeply, we can enhance our ability to interpret and predict phenotypic responses to various drugs and perturbations. Furthermore, our effort can be extended to more sophisticated methods for perturbation encoding. For example, instead of using handcrafted descriptors, we might explore embedding drug treatments using modern deep learning featurization paradigms pre-trained on thousands of compounds [65–69]. In addition to the avenues mentioned above, future work could explore using novel and powerful generative models, such as diffusion-based models, to enhance our understanding and prediction of morphological changes in response to perturbations [70]. Moreover, one could consider representing annotated genetic perturbations as numerical vectors derived from functional annotations in gene ontology [43] or antibody treatments as rich metadata arrays describing generalizable properties. Eventually, with the deployment of new high-content screening studies, we hope that future resources will allow us to test IMPA on combinations of drugs and incorporate dosage into the style conditioning process. We also believe that IMPA is compatible with additional tasks involving style transfer with context preservation, such as batch correction and denoising.

IMPA offers a valuable tool for rational experimental design in pharmaceutical research. By leveraging the model’s capabilities, researchers can conduct preliminary testing of potential drug perturbations, reducing the need for extensive experimental exploration of the chemical space. Furthermore, IMPA holds promise in optimizing molecules by assessing morphological responses, allowing for iterative *in-silico* refinement and fine-tuning of structures to achieve desired morphological changes. Additionally, IMPA can serve as a data augmentation pipeline, enabling the amplification of underrepresented perturbation classes through reliable conditional generation. These applications highlight the wide-ranging potential of IMPA in accelerating drug discovery and advancing our understanding of phenotypic responses.

## Code availability

The code for our model and link to the data is available at https://github.com/theislab/IMPA.

## Author contribution

M.L. conceived the project with contributions from F.J.T. M.L. and A.P. designed the algorithm. A.P. performed the research and implemented the algorithm. M.L. designed experiments with contributions from A.P. and F.J.T. A.P. ran all experiments. All authors contributed to the manuscript. M.L. and F.J.T. supervised the project.

## Acknowledgements

We thank Niklas Schmacke and Carlo De Donno for reviewing the paper and providing fruitful feedback. A.P. is also grateful to Leon Hetzel for his invaluable scientific mentorship and support throughout the research. M.L. acknowledges financial support from the Joachim Herz Stiftung. F.J.T. acknowledges support from the Helmholtz Association’s Initiative and Networking Fund through Helmholtz AI [ZT-I-PF-5-01]. A.P. is supported by the Helmholtz Association under the joint research school Munich School for Data Science.

## Methods

### Problem statement

Let *D* = *{*(x*_i_*, d*_i_*)*}^N^* be an image dataset where x*_i_ ∈* R*^c×h×w^* represents the i*^th^* cell image and d*_i_* = 1, …, p the index of the perturbation used to treat x*_i_*. N is the cardinality of the dataset and p is the total number of perturbation categories. Additionally, let the variable z*_i_ ∼ N* (**0**, **I**) be a random noise vector drawn from a k-dimensional multivariate Gaussian distribution and e*_j_ ∈* R*^d^*, with j = 1, …, p, be pre-computed perturbation embeddings for each treatment. We assume that the data is derived from a generative process conditioned on a content c and a perturbation-specific random style vector s *∈* R*^s^*. Specifically, we model the style encoding as a function ŝ*_i_* = f(e*_d_*, z*_i_*), where f is a linear projection of the form f: R*^d^*^+^*^k^ 1→* R*^s^*. We infer the content ĉ from the data. The embeddings e*_di_* can be either fixed and obtained from a prior representation of the treatments or randomly initialized and trained.

Our goal is to approximate the unknown conditional probability distribution p*^∗^*(x*|*c, s) through a parametrized generative model p*_θ_*(x*|*c, s) that can be actively sampled. In this framework, the latent representations c and s respectively control the visual content of the generated sample and its perturbation-specific style. In the ideal case where our learned p*_θ_* converges to p*^∗^*, any sample x^*_i_* from p*_θ_*(.*|*ĉ*_i_*, ŝ*_i_*) appears like real examples of cells treated with perturbation d*_i_*, but conserving visual features encoded by ĉ*_i_*.

Under such a formulation, our model is capable of performing transformation of data point x*_i_*, coming from the real d*_i_^th^* perturbation modality, to an inferred x^*^′^* image perturbed by condition *^′^ d*= d*_i_*. This is achieved by sampling from p*_d_*(.*|*ĉ*_i_*, ŝ*^′^*), where ŝ*^′^* is the style derived for d*^′^*.

Specifically, x^*^′^* represents what the input x*_i_* would look like under a different perturbation type.

In what follows, we describe the implementation and training process of IMage Perturbation Autoencoder (IMPA), a generative adversarial model performing phenotypic transformations on cell perturbation images.

### Training

We develop our model as a conditional generative adversarial network (cGAN) for image-to-image translation. The architecture and underlying working principles are inspired by StarGANv2 [38]. The main backbone of the architecture is constituted by an image encoder E, a decoder D, a discriminator Dis, a style encoder E*_sty_* and a perturbation encoder f.

During training, the input tuple (x*_i_*, d*_i_*) is fed to the model. The image x*_i_* is used to infer the content ĉ*_i_* as follows:

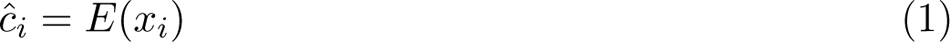

where E models the content space. To teach the model how to induce a distribution shift on the images to a different condition, we perform input transformation during training. More specifically, for each training pass, a condition d*^′^ d*= d*_i_* is sampled from a discrete uniform distribution over indices *I − {*d*_i_}*, with *I* = *{*1, …, p*}*, and used to select the embedding e*_d_′* of the corresponding perturbation. IMPA is a probabilistic generative model. To induce a non-deterministic between-style transition, we concatenate e*_d_′* with a random noise vector z*_i_* and encode such combination to a style vector via f as:

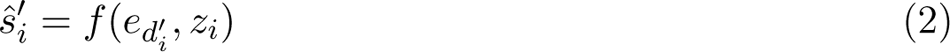

The derived style is used to condition the generative model function approximated by the decoder D. In this regard, the style vector ŝ*^′^* is meant to induce the generator to decode ĉ*_i_* into an image x^*^′^* with characteristics typical of perturbation d*^′^*. For brevity, we refer to the combination between E and D as *generator* and indicate it with G. Specifically, we indicate the succession of encoding and decoding passes as G(x, s) = D(E(x), s)

To favour realism in the synthesized image, we couple the described autoencoder architecture with a multi-task discriminator Dis. Provided an input, the discriminator yields p different probability values, one for each perturbation class. Every output unit j performs the task of predicting the probability that the discriminator’s input is real within domain j. Consequently, in our model, a generated image can be equally realistic for multiple domains which are not mutually exclusive. Based on the generator-discriminator interplay, we define the adversarial loss as:

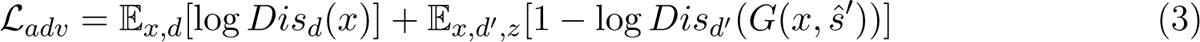

where Dis*_d_*(x) stands for the d*^th^* output of the multi-task discriminator. Equation 3 essentially describes how the generator and the discriminator are jointly trained. The discriminator is tweaked to maximize the function *L_adv_* by learning to predict an observation x*_i_* as real in its domain d*_i_*. Conversely, the generator strives to deceive its adversary by maximizing the probability that its generated output conditioned on ŝ*^′^* is deemed as a real input from domain d*^′^*.

The problem we are solving is a typical instance of distribution alignment, where the conditional distribution learned by the generator is matched to the real data distribution through a minimax optimization problem. To further push the generator network to associate an image to its domain-specific style, we train a style encoder E*_sty_*: R*^c×h×w^ 1→* R*^s^* to approximate the style vector ŝ*^′^* starting from a transformed image x^*^′^*. This is accomplished through the following loss function:

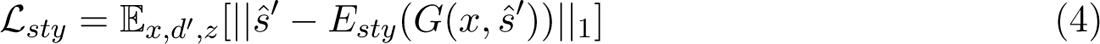

variant characteristics of the input image, we push the model to reconstruct the original cell from its transformed counterpart. To this end, we condition the decoding of x^*^′^* on the style vector E*_sty_*(x*_i_*) to retrieve an approximation x^*_i_* of x*_i_*. Such a process further ensures that the style space learned by f is in line with that implemented by E*_sty_* and they cause a compatible shift in the generated distribution. Let x^*^′^* = G(x, ŝ*^′^*), we define the cycle consistency loss [46] as follows:

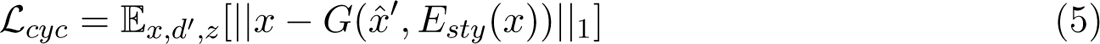

Finally, following prior work, we enforce diversity in the generated output by a style diversification loss [71]. Given two independent random noise vectors z^(1)^ and z^(2)^ from the same distribution and the respective styles ŝ*^′^*^(1)^ = f(d*^′^*, z^(1)^) and ŝ*^′^*^(2)^ = f(d*^′^*, z^(2)^) for the same perturbation d*^′^*, we maximize the heterogeneity between their generated outputs through the following style diversification loss:

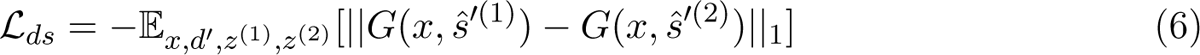

Equation 6 ensures that the random vector is not ignored and a single perturbation can produce a distribution of diverse counterfactual responses.

Collecting all the terms together, the generator is trained to minimize the final loss:

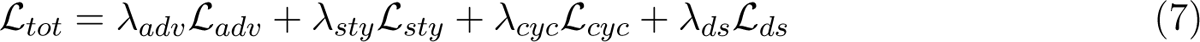

where the λ coefficients delineate scale parameters to define the relative importance of each component. All scaling terms are kept constant during training except for λ*_ds_* which is decayed linearly across the budget of iterations until it reaches a value of 0.

Having depicted the optimization objective of our model, each training iteration proceeds through the following steps:

1. An input (x*_i_*, d*_i_*) is sampled
2. A perturbation d*^′^ d*= d*_i_* and two random vectors z^(1)^ and z^(2)^ are sampled from the respective distributions
3. The styles ŝ*_i_* = f(e*_d_′*, z*_i_*) and ŝ*_i_* = f(e*_d_′*, z*_i_*) are inferred and used to generate transformed inputs to approximate Equation 6
4. The generated output x^*^′^*= G(x*_i_*, ŝ*^′^*^(1)^) is used to approximate Equation 3 together with the real batch
5. E*_sty_* computes style vectors from x*_i_* and x^*^′^* = G(x*_i_*, ŝ*^′^*^(1)^) to approximate Equations 4 and
6. *L_tot_* is calculated as described in Equation 7 and minimized via gradient descent

### Testing

Given an input observation (x*_i_*, d*_i_*) and a perturbation d*^′^* for which we would like to generate a prediction, IMPA addresses the question: *what morphological changes would be induced in* x*_i_ had it been perturbed by* d*^′^ instead of* d*_i_*? Provided the components described in the previous section, testing is carried out through the following steps:

1. Sample z*_i_ ∼ N* (**0**, **I**) and collect e*_d_′*
2. Compute the style ŝ*^′^* = f(e*_d_′*, z*_i_*)
3. Infer the content ĉ*_i_* = E(x*_i_*)
4. Perform decoding on ĉ*_i_* conditioned on the style vector ŝ*^′^* to obtain the result of the prediction x^*^′^* = D(ĉ*_i_*, ŝ*^′^*)

### Architecture

#### Normalization method and style conditioning Instance Normalization

Instance normalization (IN) [72] differs from batch normalization (BN) [73] in that it produces observation-specific scales and shifts instead of learning them for the whole batch. Given an observation x*_tchw_*, where t indexes the batch dimension, c the channel, h the height and w the width, IN standardizes an observation as follows:

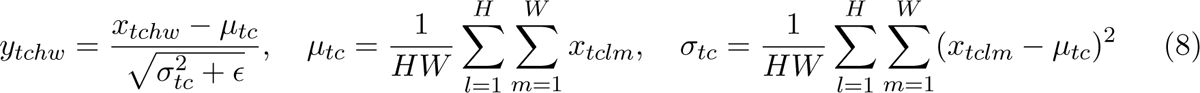

where µ*_tc_* and σ*_tc_* are the pixel mean and variance of a single instance computed across spatial dimensions. H and W indicate the height and the width of the images, whereas y*_tchw_* is the outcome of the normalization. Since IN is implemented as a differentiable neural network layer, additional scaling and shifting parameters γ and β are learned during training, giving rise to the following expression:

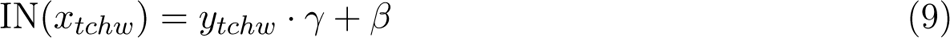

Notably, Equation 9 implements an affine transformation of the input.

### Adaptive Instance Normalization (AdaIN)

IN has been demonstrated to normalize the input to a specific style controlled by learned affine transformation parameters. Intuitively, if we can learn to shift and scale the convolutional feature space of an observation based on a class-specific style, then we can implement a generator performing style transfer on demand. Thus, we employ ADAptive INstance Normalization (AdaIN) [33] to perform domain translation in the decoder of our model.

Given an input x and a style y, AdaIN computes the following transformation:

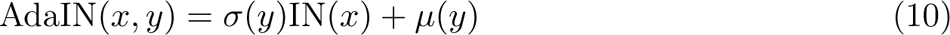

where σ(y) and µ(y) are learned style-dependent affine parameters that normalize the image x to style y. By operating in the convolutional feature space, AdaIN facilitates domain transfer by enhancing the feature channels responsible for conveying a determined visual style response.

### Residual block

All the image-processing components in our model are implemented as a residual block [74]. Any input x is simultaneously fed to a convolutional stack called *residual mapping* and a shallow *skip connection*. Subsequently, the results from said branches are summed to produce the residual block output.

An overview of the network layers in the residual block can be observed in Table 1. Encoder and decoder residual blocks differ in the way they modify the dimensionality in the spatial dimension. More precisely, the encoder network reduces the image height and width by a factor of two through average pooling, whereas the decoder upsamples feature maps spatially via nearestneighbor interpolation. Since the residual mapping produces changes in dimensionality, the skip connection is equipped with average pooling or upsampling layers and a 1*×*1 convolution to match the spatial and depth dimensions of the residual branch.

**Table 1.**
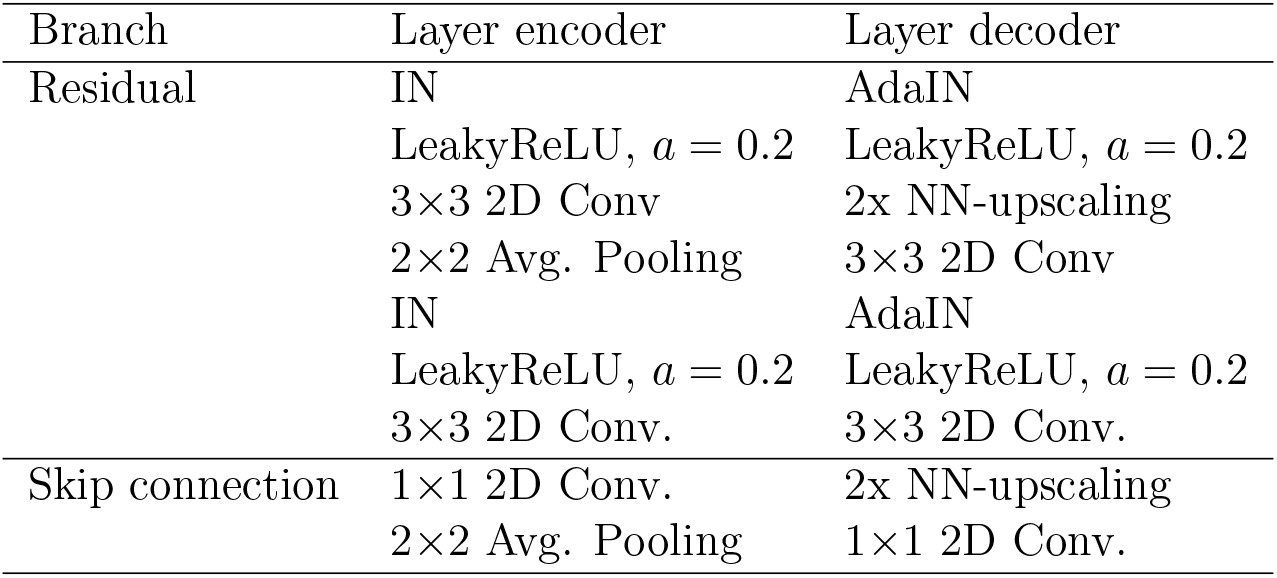
The list of layers composing the residual architecture. The layers are placed in the same order as they are found in the model. Encoder and decoder differ by the normalization method, the ordering and the dimensionality-altering layer. The encoder uses instance normalization and downsamples spatially via average pooling. The decoder is conditioned on the style through AdaIN and upsamples its input through the Nearest Neighbor (NN) interpolation.

### Generator

The architecture of the generator is illustrated in Table 2. It consists of an encoder and a decoder that implement the image translation task. The encoder computes the latent content starting from an image. Such a representation is highly dimensional (512 *×* 12 *×* 12) to allow for the preservation of spatial features. Notably, the decoder network mirrors the encoder as shown in Table 1. While the encoder does not receive any information on the perturbation style, the decoder is conditioned on it via AdaIN. More formally, the convolutional features of the decoder residual block are scaled based on the perturbation style vector. To achieve perturbation-specific scaling of the residual blocks using AdaIN, the style vector ŝ *∈* R*^s^* is passed through a tunable linear layer that approximates the functions σ(ŝ) and µ(ŝ) to compute the affine transformation reported in Equation 10.

**Table 2.** The list of layers constituting the generator network. The entries down. and up. ResBlock derive from the modular structure described in Table 1, with, respectively, average pooling (encoder) or NN interpolation (decoder) layers. Their dimension-preserving counterpart is indicated as ResBlock.

| Module | Layer | Dimension |
| --- | --- | --- |
| Encoder | 3×3 2D Conv. | 64×96×96 |
|  | Down. ResBlock | 128×48×48 |
|  | Down. ResBlock | 256×24×24 |
|  | Down. ResBlock | 512×12×12 |
|  | ResBlock | 512×12×12 |
|  | ResBlock | 512×12×12 |
| Decoder | ResBlock with AdaIN | 512×12×12 |
|  | ResBlock with AdaIN | 512×12×12 |
|  | Up. ResBlock with AdaIN | 256×24×24 |
|  | Up. ResBlock with AdaIN | 128×48×48 |
|  | Up. ResBlock with AdaIN | 64×96×96 |
|  | Instance norm. | 64×96×96 |
|  | LeakyReLU | 64×96×96 |
|  | 1×1 2D Conv. | 3×96×96 |

### Discriminator and style encoder

The architectures of the discriminator and style encoder are illustrated in Table 3. The discriminator is a convolutional neural network consisting of downsampling residual blocks and a final 2D convolutional layer with 3 *×* 3 kernel. Let p be the number of available perturbations and x an input set with batch size T, number of channels C, width W and height H. The discriminator acts on the input reducing it to an output array shaped T *×* p *×* 1 *×* 1. Successively, the spatial dimensions are trimmed and a sigmoid activation is applied to each node. The resulting T *×* p tensor represents the class-specific predictions for the input. The inferred style encoder has the same organization as the discriminator network. However, it projects an image onto a T *×* s tensor, where s is the dimensionality of the style space. No activation function is applied to the output of the module.

### Perturbation encoder

The perturbation encoder carries out a simple linear projection from the perturbation embedding to the approximated style space. Therefore, it is implemented as a simple fully-connected layer with no activation.

**Table 3.** The discriminator and style encoder architectures implemented by IMPA. With p, we indicate the number of perturbation categories, whereas s is the dimensionality of the style space.

| Module | Layer | Dimension discr. | Dimension style enc. |
| --- | --- | --- | --- |
| Discriminator | 3×3 2D Conv. | 64×96×96 | 64×96×96 |
|  | Down. ResBlock | 128×48×48 | 128×48×48 |
|  | Down. ResBlock | 256×24×24 | 256×24×24 |
|  | Down. ResBlock | 512×12×12 | 512×12×12 |
|  | Down. ResBlock | 512×6×6 | 512×6×6 |
|  | Down. ResBlock | 512×3×3 | 512×3×3 |
| | 3×3 2D Conv. | $p \times 1 \times 1$ | $s \times 1 \times 1$ |
| | Reshape | $p$ | $s$ |
| | Sigmoid | $p$ | - |

### Additional aspects Gradient penalty

We control the magnitude of the discriminator’s updates by adding a gradient penalty term *L_reg_* with coefficient λ*_reg_* to the loss. *L_reg_* computes the sum of squared gradients of the discriminator’s output with respect to all the pixels of the real input on which it is trained. The sum of squared gradients is then averaged across the batch dimension as follows:

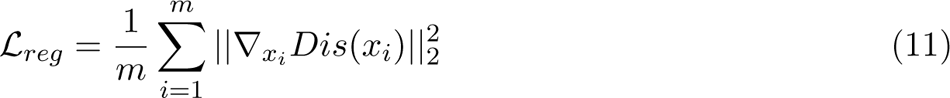

where x*_i_* is the i*^th^* image of a real batch with m samples and *∇_x_* the gradient of the discriminator’s output with respect to the pixels of the input image x*_i_*.

### Data augmentation

Each training image undergoes random vertical and horizontal flips with a probability of 0.3 before training as a form of data augmentation. Moreover, we add random noise elementwise to all images before feeding them to the encoder.

### Balanced sampling

When the training set is unbalanced, we observe mode collapse on perturbation classes consisting of a small number of examples. More specifically, the generator learns to ignore the content space from the encoder network and transforms every input into the same prediction regardless of its original spatial features. We, therefore, introduce balanced sampling, where the probability of including an observation is decreased proportionally to the frequency of its perturbation class. Thereby, too ubiquitous categories are precluded from over-populating training iterations.

### Weight initialization

To prevent the arising of vanishing and exploding gradients in the context of non-linear activation functions, we adopt the popular *He weight initialization* [75]. This method is specifically designed to stabilize training with the ReLU activation function and its variants by controlling the weight variance upon initialization. In practical terms, the He approach draws the initial weights w*_l_* for layer l from the following distribution:

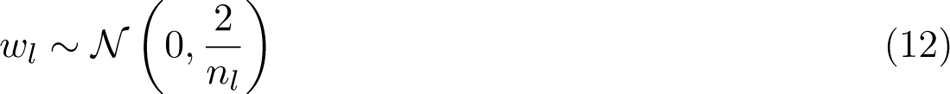

where n*_l_* is the number of input neurons of l. To fulfil the requirement of having zero-centred weights and outputs, the bias is set to 0 at the beginning.

### Training evaluation

During training, we perform transformations of test set data points to random target drugs and evaluate the average Fréchet Inception Distance [57] distance between the translated results and the real perturbation images from the target class. GAN evaluation is an ongoing field of research and existing metrics do not always reflect the visual quality of the model. Moreover, the probabilistic behaviour of the model complicates the reliability of its evaluation by adding a layer of uncertainty. Therefore, model selection is carried out by combining metric calculation, loss convergence and qualitative evaluation of sample transformations. In cases of comparable performance, the most parsimonious model in terms of resources was selected.

### Baselines StarGANv1

StarGANv1 [53] performs conditional image-to-image translation across multiple domains using a single adversarial network. Similar to IMPA, the architecture is based on an autoencoder model that maps input data to a latent space shared among domains. During training on a data point, a different perturbation is sampled as a one-hot encoded vector. Said vector is used to couple decoding by broadcasting and appending it to the input image as additional feature maps. Subsequently, the decoded output is submitted to a multi-task discriminator that tries to simultaneously predict whether the result is real or fake and to which class it belongs. By learning to mislead the discriminator, the autoencoder acts as a generator of transformed images. As the generation is fully conditioned on a lookup table for the domain labels, the model is deterministic.

#### DMIT

Disentanglement for Multi-mapping Image-to-Image Translation (DMIT) [55] performs multimodal and multi-domain image transformation via content and style disentanglement. It assumes that each image in the dataset is disentangled across three latent representations referred to as content, style and label spaces. Specifically, the image content and style are inferred through encoder networks, whereas the label is one-hot encoded and used for conditioning a residual decoder. The training was divided into a disentanglement and a translation path. The former pushes the model to reconstruct the original input from disentangled spaces and regularizes the style encoding to a multivariate normal distribution. Conversely, the translation path implements the adversarial mechanism where styles and domains are swapped and a domain-conditioned discriminator is deceived into mislabeling generated images as real. What distinguishes DMIT from the other models is the presence of two latent regression terms that try to predict the inferred content and style vectors from a generated image.

#### DRIT++

DRIT++ [54] is a popular architecture in the field of domain adaptation. It encodes an image into two disentangled spaces: a domain-invariant content encoding and a condition-specific attribute vector. During training, random pairs of images from different domains are sampled and their attributes are swapped in the decoding process. From the decoded outputs, a multi-task generator tries to predict if the synthesized images are real and to what domain they belong. To force the content space to be completely class-agnostic, an additional content discriminator is deceived into failing to predict the domain of origin from the content vector. Moreover, a cross-cycle consistency loss is computed by training the model to invert the style exchange process. All steps are conditioned on one-hot domain encodings, which allow guiding the model to perform translation to a given class during inference.

### Mol2Image

Mol2Image [26] is a conditional generative model originally applied to the task of cell image generation. The authors implemented a multiscale version of Glow [76], a generative framework based on normalizing flows. Cell images are encoded to multiple levels of latent representations at different scales obtained by decomposing the input images into coarse and fine-grained down-sampled representations via a Haar wavelet image pyramid. The latent codes are regularized to a Gaussian distribution whose parameters are conditioned on the next level of the pyramid and a dense drug representation is obtained via Graph Neural Networks (GNNs). To further enhance drug conditioning, the authors train the model with a contrastive loss, which pulls the latent representations of images closer to the embedding of the drug used to perturb them. In the generation phase, training and unseen drugs are used to condition the sampling of latent codes for image synthesis, therefore providing a tool to infer the morphological effects of drug perturbations on cell data.

### Untransformed images

Alongside neural network baselines, we report evaluation scores computed on untransformed control images. In other words, we compare the source images before translation with the true perturbed examples from the dataset. This provides a lower bound on the model performance as it represents the evaluation score we would expect if the network could not translate the inputs at all.

### Evaluation metrics

#### GAN evaluation metrics

##### Fréchet Inception Distance (FID)

The Fréchet Inception Distance (FID) [57] measures the difference between the real and generated image distributions in the feature space learned by an Inception V3 model [63] pre-trained on the ImageNet dataset [77]. Batches of true and fake samples are passed through the model and 2048-dimensional encodings are extracted from the last pooling layer. This produces visually-relevant feature vectors that are used as a proxy to define the perceptual distance between the compared distributions. Specifically, for both real and generated batches, the mean and the covariance of the derived encodings are calculated. Let m*_r_* and m*_g_* be the mean Inception V3 embedding vectors for real and generated images and C*_r_* and C*_g_* be the relative covariance matrices. We want to compare the feature distribution heterogeneity between true and fake images through the Frechét distance metric. To this end, we assume that both real and generated inception features are distributed as multivariate normal distributions and that their means and the variances can be used to compute the Frechét distance between them. As a result, the FID is expressed as:

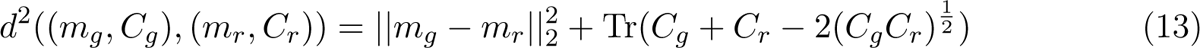

where Tr refers to the trace of a matrix and *||*.*||*^2^ is the squared L^2^-norm. To make it comparable with distance measures, we report the inverse of the FID score.

##### Density and coverage

FID produces a summary of the distance between two distributions and does not separately evaluate the fidelity and diversity of generated sample to the real data [58]. As a result, we introduce two additional metrics to assess our generative model: Density and Coverage.

Let X and Y be respectively samples of real and generated images. We approximate a manifold in X as:

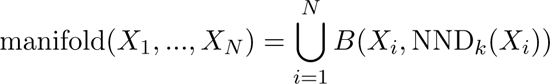

where B(x, d) is the sphere defined around the point x with radius d and NND*_k_*(x) is the distance of the furthest nearest neighbor from x in a neighborhood of size k. Given a manifold around the true data X, we measure density:

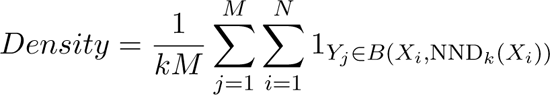

More in detail, density measures the proportion of real neighborhoods containing generated samples Y*_j_* across all fake examples. This process yields high values in regions with a high density of real samples and avoids overestimating the manifold around outliers.

Conversely, coverage measures the extent to which generated samples recover the whole real sample space and is defined as:

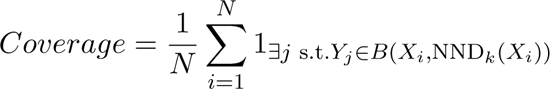

Namely, coverage is higher when a large number of real observations have a fake example in their neighborhood.

For both measures, higher values indicate better generation quality.

##### Accuracy of MoA classification

We additionally evaluate model predictions for drug perturbations based on the frequency with which they are correctly labeled with their actual mode of action by a pre-trained classifier. More specifically, we train a classifier with the same architecture as the model discriminator (see **Architecture**) to predict MoA labels on the true images. Subsequently, we transform control cells into their drug-perturbed counterparts and infer the MoA on the generated images using our classifier. The higher the accuracy with which a model’s prediction is labeled with its true MoA, the better the generation output.

##### Evaluation on morphological features

We used the CellProfiler 4.2.1 [51] software to calculate reliable morphological features of both generated and real images from the test set for comparison. Through this, we aim to evaluate if the transformations produced by our model on control images reproduce the phenotypic shift caused by perturbation. A custom CellProfiler script is executed on a dedicated graphical user interface to perform segmentation and extract phenotypic profiles for cells. Briefly stated, the software identifies cell nuclei via Otsu thresholding [78] and uses them as seeds to outline the corresponding objects in the cytoplasm channels via Watershed segmentation [79]. The outcome of this step is used to calculate an array of features quantifying cell shape, size, intensity distribution and texture across high-content channels. In total, we obtain 356 morphological features per image.

### Datasets

#### BBBC021

We choose the BBBC021v1 dataset [80] from the Broad Bioimage Benchmark Collection. It contains images from a high-content screening assay featuring p53 wild-type MCF-7 cell lines of breast cancer. In the experiment, each well is fluoresced across three channels: nucleus (stained with DAPI), β-tubulin (marked with an anti-β-tubulin antibody) and F-actin (stained with Phalloidin). Precisely, the assay includes a series of 55 96-well plates grouped into batches acquired over ten weeks. Each plate contains six negative controls treated with dimethyl sulfoxide (DMSO) only and six positive controls incubated with taxol. The remaining wells are exposed to active compounds in triplicates and at 8 different concentrations for 24 hours. The original dataset reports the screening of 113 molecules. However, we choose to utilize the version of the dataset annotated by [47], where only 38 perturbations are reported based on their potential to induce a phenotype at the used concentrations. In general, 12 MoAs were annotated on the compounds. From this group, the authors claim that only 6 MoAs were distinguishable in the images, whereas the rest were mined from the literature. The dataset consists of 2528 full-well images with a resolution of 1024*×*1280 pixels each.

#### BBBC025

The BBBC025 dataset [81] includes images of U2OS cell lines for which gene knockdown was performed using short-hairpin RNA (shRNA) interference. In this study, a selection of 41 genes actively expressed in U2OS cells was targeted by synthetic 6-nucleotide-seed shRNA molecules. For every gene, six or more shRNA sequences were selected for a total of 350 different perturbations. Cells were cultured in 384-well plates. Each plate configuration was replicated six times, with additional negative (untreated) and positive (treated with puromycin) controls to assess infection efficiency. Upon 96 hours of incubation, images were acquired through the Cell Painting assay [82]. Specifically, eight cellular sub-structures were marked separately by five different fluorescent compounds and imaged at 20x magnification. After post-processing, 3072 full-well 1084*×*1084 images were collected and deployed.

#### RxRx1

The RxRx1 dataset [83] involves 16-bit fluorescence microscopy images of cell lines perturbed by 1,138 distinct small interfering RNAs (siRNA). Similarly to BBBC025, fluorescent images were obtained through the Cell Painting assay across six channels highlighting heterogeneous cellular compartments. Four different cell lines (HUVEC, RPE, HepG2, and U2OS) were analyzed in distinct numbers of batches grouping 384-well plates. In total, 51 experimental batches were produced, with HUVEC being the most frequent cell line. In this framework, randomized treatments were plated in combination with 30 controls and one non-treated well. Finally, the recovered images were downsampled to resolution 512*×*512 pixels to support downstream applications. Unlike the BBBC025 dataset, RxRx1 is not annotated with the names of the silenced genes.

### Data pre-processing

#### Illumination correction algorithm and normalization

To reduce illumination artefacts due to plate effects, we perform the illumination correction algorithm described in [84] and revisited in [14] on full-resolution images. First, images from each plate are aggregated and used to compute a flatfield image by taking the pixelwise 10th percentile across each plate group. Successively, the result is smoothed out by applying a Gaussian filter with a standard deviation of 50. The pixels of each well are divided by the flatfield image of the respective plate to yield the background intensity distribution for adjustment. Furthermore, the dynamic range of each image is enhanced by clipping the pixel values at 1 and computing the natural logarithm of the intensities. Following the original implementation, we additionally clip the transformed pixel intensities at a maximum of 5 to avoid outliers. The output is linearly re-scaled between 0 and 255 and quantized to 8 bits.

#### BBBC021

The full-resolution images were corrected for illumination and cropped into patches centred around single nuclei in squares of 96*×*96 pixels. To this end, we used the coordinates of 454,793 cells pre-computed by [47] and excluded objects at less than 48 pixels from the image borders.

Out of the 38 drugs reported in the dataset, 4 were ruled out due to undisclosed chemical structures. To remove corrupted images displaying empty wells or pixel noise, we empirically filtered out all the images with a pixel variance lower than 2000. Moreover, we reduced the number of highly blurred images in the dataset by computing a normalized perceptual blur metric [85] over each crop, where 1 indicates maximal blur and 0 absence thereof. A threshold for acceptance of an image was set at 0.65 to avoid over-pruning while still filtering excessively low-quality wells. Control images tend to produce a much larger number of cells. As a result, we downsampled the number of DMSO-treated cells to 150 random controls per plate to favour balanced batches during training. Alongside the complete data, we created a reduced version of the dataset for comparison with the baseline image-to-image translation architectures. Specifically, we selected Cytochalasin B, Taxol, Vincristine, AZ258 and AZ138 inducing actin disruption, tubulin destabilization, tubulin stabilization, Aurora kinase inhibition and Eg5 inhibition, respectively. The drug for each mode of action was manually selected based on image quality and effect size. We additionally kept DMSO as a perturbation category, yielding a 6-class dataset. As a result, the full dataset contains 97,504 observations, whereas its subset version spans 20,313 well patches. Due to class imbalance and scarcity of remaining data points for some of the drug conditions, we decided to dedicate 90% of the dataset to training and 10% to testing.

#### BBBC025

After illumination correction, the nuclear channels from BBBC025 were thresholded via the Otsu method [78]. Cell centres were estimated based on the object mask and used to create 96*×*96 patches around the cells. Moreover, the dataset size was controlled by only keeping 300 cells from each shRNA condition after removing acquisition noise with a variance threshold of 2000. The shRNA molecules, representing a perturbation condition each, were distinguished based on their target gene, resulting in a total of 42 categories (including the negative controls). Thus, we recovered a dataset containing 105,300 data points and split it into 80% training and 20% test subsets.

#### RxRx1

In analogy to BBBC025, the images of RxRx1 were corrected for illumination effects within the plates and their nuclear channels were segmented via Otsu thresholding [78]. We derived patches of dimension 96*×*96, including one or a few cells depending on the bulkiness of the well. For simplicity, we focused on the U2OS cell line of osteosarcoma as our model does not account for inter-cell line transformations. A maximum of 100 cell images per well was kept assuming constant effects across the field of view. In total, we obtained more than 170k images for 1,139 conditions (1,108 treatments, 30 control treatments and 1 untreated example per plate). We further subset the conditions to the 1,070 perturbations whose target genes are included in the Gene2Vec model. Finally, we partitioned the dataset into 80% training and 20% test set.

### Perturbation embedding

#### Drug compounds in BBBC021

As drug embeddings in BBBC021, we employ the 2D chemical features computed by the RDKit Python package [39]. Every compound with a disclosed chemical structure can be associated with a 201-dimensional descriptor vector computed directly from its SMILES representation. The feature vectors were additionally standardized to account for scale heterogeneity and filtered out for poorly variable dimensions with standard deviation under 0.01 across drugs [41]. The recovered embeddings include 160 descriptors for each available drug. To preserve their physiochemical meaning, the drug embeddings are made non-trainable.

#### shRNAs and siRNAs in BBBC025 and RxRx1

The siRNA and shRNA target genes were embedded using the Gene2Vec model [40]. Gene2Vec is a computational method inspired by Word2Vec [86] that learns distributed representations of genes in a vector space. By capturing the semantic and functional relationships between genes, Gene2Vec generates 200-dimensional fixed-length vectors that encode co-expression patterns in transcriptomics datasets.

#### Hyperparameters

**Table 4.**
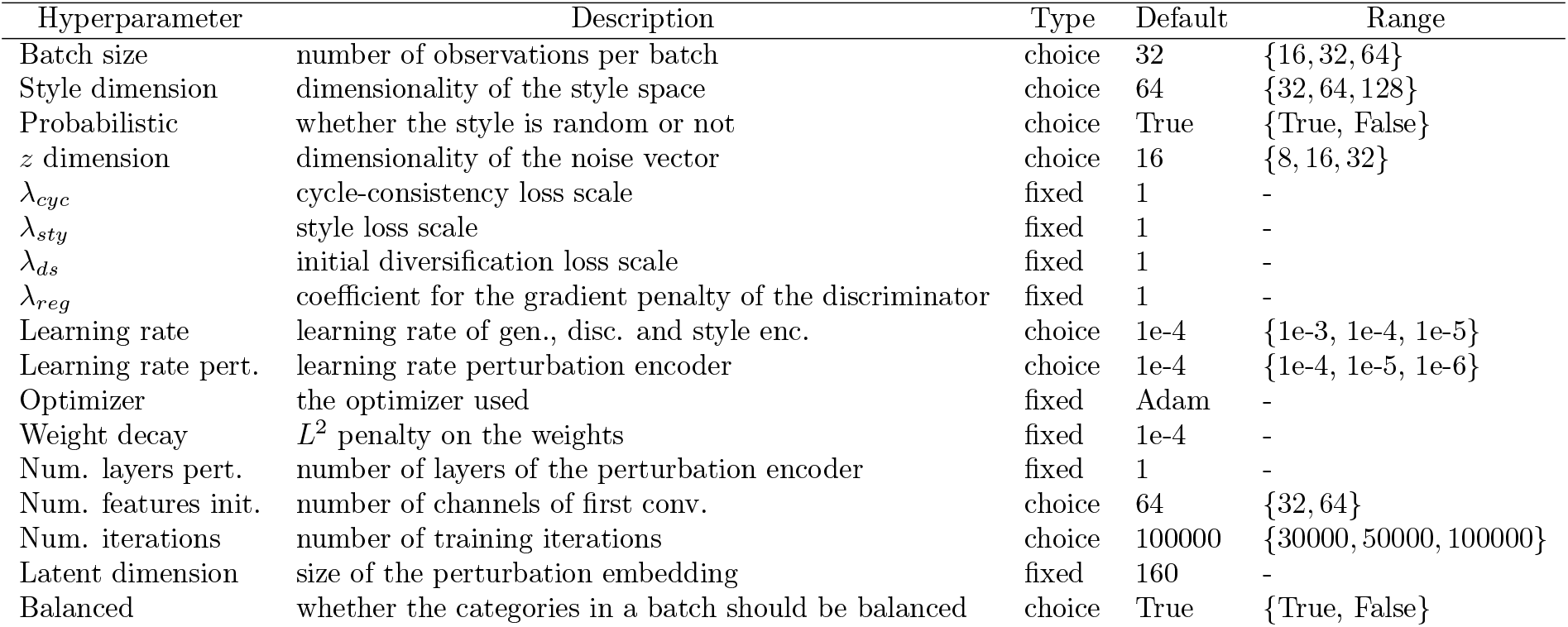
The tunable hyperparameters to train IMPA on the BBBC021 subset with 5 drugs. The column *Type* defines whether the parameters were fixed as in [38] or optimized via randomized grid search. Conversely, *Range* refers to the list of options explored during the tuning. The *Default* column indicates the value in *Range* that was selected after optimization.

| Hyperparameter | Description | Type | Default | Range |
| --- | --- | --- | --- | --- |
| Batch size | number of observations per batch | choice | 32 | {16, 32, 64} |
| Style dimension | dimensionality of the style space | choice | 64 | {32, 64, 128} |
| Probabilistic | whether the style is random or not | choice | True | {True, False} |
| $z$ dimension | dimensionality of the noise vector | choice | 16 | {8, 16, 32} |
| $\lambda_{cyc}$ | cycle-consistency loss scale | fixed | 1 | - |
| $\lambda_{sty}$ | style loss scale | fixed | 1 | - |
| $\lambda_{ds}$ | initial diversification loss scale | fixed | 1 | - |
| $\lambda_{reg}$ | coefficient for the gradient penalty of the discriminator | fixed | 1 | - |
| Learning rate | learning rate of gen., disc. and style enc. | choice | 1e-4 | {1e-3, 1e-4, 1e-5} |
| Learning rate pert. | learning rate perturbation encoder | choice | 1e-4 | {1e-4, 1e-5, 1e-6} |
| Optimizer | the optimizer used | fixed | Adam | - |
| Weight decay | $L^2$ penalty on the weights | fixed | 1e-4 | - |
| Num. layers pert. | number of layers of the perturbation encoder | fixed | 1 | - |
| Num. features init. | number of channels of first conv. | choice | 64 | {32, 64} |
| Num. iterations | number of training iterations | choice | 100000 | {30000, 50000, 100000} |
| Latent dimension | size of the perturbation embedding | fixed | 160 | - |
| Balanced | whether the categories in a batch should be balanced | choice | True | {True, False} |

**Table 5.** The hyperparameters used to optimize IMPA on the BBBC021 dataset with all 34 drugs. A description of the hyperparameters is provided in Table 4.

| Hyperparameter | Type | Default | Range |
| --- | --- | --- | --- |
| Batch size | fixed | 32 | - |
| Style dimension | choice | 64 | {8, 16, 32, 64} |
| Probabilistic | choice | True | {True, False} |
| $z$ dimension | choice | 64 | {8, 16, 32, 64} |
| $\lambda_{cyc}$ | fixed | 1 | - |
| $\lambda_{sty}$ | fixed | 1 | - |
| $\lambda_{ds}$ | fixed | 1 | - |
| $\lambda_{reg}$ | fixed | 1 | - |
| Learning rate | fixed | 1e-4 | - |
| Learning rate pert. | fixed | 1e-4 | - |
| Optimizer | fixed | Adam | - |
| Weight decay | fixed | 1e-4 | - |
| Num. layers pert. | fixed | 1 | - |
| Num. features init | fixed | 64 | - |
| Num. iterations | fixed | 100000 | - |
| Latent dimension | fixed | 160 | - |
| Balanced | choice | True | {True, False} |

**Table 6.** The hyperparameters used to optimize IMPA on the BBBC025 dataset. A description of the hyperparameters is provided in Table 4.

| Hyperparameter | Type | Default | Range |
| --- | --- | --- | --- |
| Batch size | fixed | 32 | - |
| Style dimension | fixed | 64 | - |
| Probabilistic | fixed | True | - |
| $z$ dimension | choice | 32 | {8, 16, 32, 64} |
| $\lambda_{reg}$ | fixed | 1 | - |
| $\lambda_{cyc}$ | fixed | 1 | - |
| $\lambda_{sty}$ | fixed | 1 | - |
| $\lambda_{ds}$ | fixed | 1 | - |
| Learning rate | fixed | 1e-4 | - |
| Learning rate pert. | choice | 1e-4 | {1e-6, 1e-5, 1e-4} |
| Optimizer | fixed | Adam | - |
| Weight decay | fixed | 1e-4 | - |
| Num. layers pert. | fixed | 1 | - |
| Num. features init | fixed | 64 | - |
| Num. iterations | fixed | 100000 | - |
| Latent dimension | choice | 100 | {50, 100, 150} |
| Balanced | fixed | True | - |

**Table 7.** The proposed tunable hyperparameters to train IMPA on the RxRx1 datasets. A description of the hyperparameters is provided in Table 4.

| Hyperparameter | Type | Default | Range |
| --- | --- | --- | --- |
| Batch size | fixed | 32 | - |
| Style dimension | fixed | 64 | - |
| Probabilistic | fixed | True | - |
| $z$ dimension | choice | 16 | {8, 16, 32, 64} |
| $\lambda_{reg}$ | fixed | 1 | - |
| $\lambda_{cyc}$ | fixed | 1 | - |
| $\lambda_{sty}$ | fixed | 1 | - |
| $\lambda_{ds}$ | fixed | 1 | - |
| Learning rate | fixed | 1e-4 | - |
| Learning rate pert. | choice | 1e-4 | {1e-6, 1e-5, 1e-4} |
| Optimizer | fixed | Adam | - |
| Weight decay | fixed | 1e-4 | - |
| Num. layers pert. | fixed | 1 | - |
| Num. features init | fixed | 64 | - |
| Num. iterations | fixed | 100000 | - |
| Latent dimension | choice | 100 | {50, 100, 150} |
| Balanced | fixed | True | - |

#### Software and computational resources

IMPA is fully implemented in Python 3.9.7. For deep learning models, we used PyTorch 1.8.0 with torchvision 0.9.0.

All experiments were run on different GPU servers with the following characteristics:

- 2x Intel(R) Xeon(R) Platinum 8280L CPU with 28 cores / 2.70GHz / 16x Tesla V100 GPUs with 32GB of RAM per card
- 2x Intel(R) Xeon(R) 6230 CPU with 20 cores / 2.1GHz / 2x Tesla V100 GPUs with 16GB of RAM per card
- 2x AMD EPYC 7742 CPU with 256 cores / 2250.0000 MHz / 8x A100-SXM4 GPU with 40GB of RAM per card

---

**Supplementary Figure 1.**
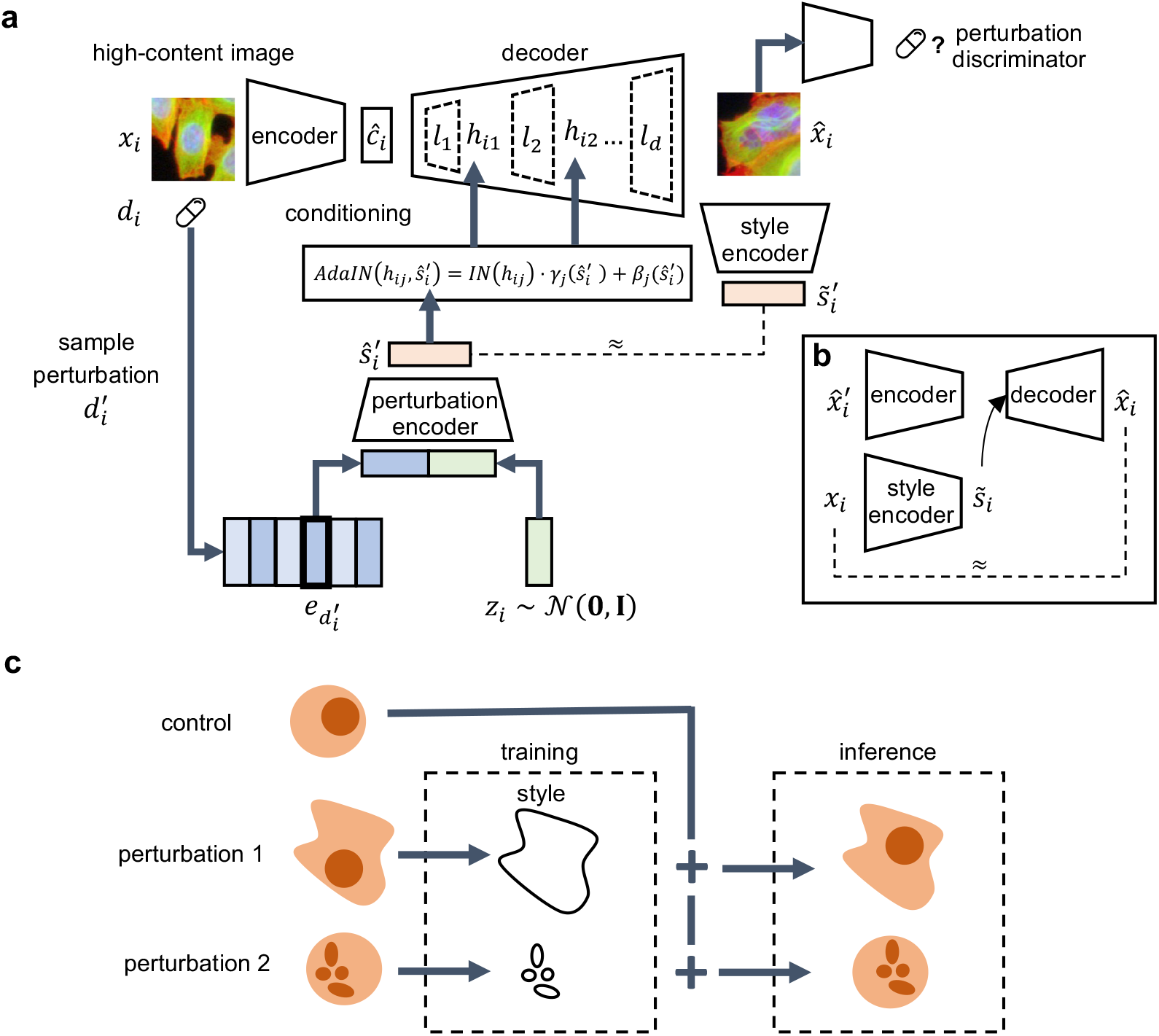
Model architecture and working principles of IMPA. (a) A visual description of the full architecture of IMPA. A control cell image x*_i_* is encoded into a content representation. When applying transformations to the input, a dense embedding of the target perturbation is collected and concatenated with a normally-distributed random vector. A low-dimensional projection of the concatenation constitutes the style space which is used to condition every layer l*_i_* of the decoder via the AdaIN method [33]. With h*_ij_* we indicate the output of the j*^th^* decoder layer on image i. The transformed output is used to induce a discriminator net into predicting that the decoded image comes from the target perturbation batch. Moreover, a style encoder is trained to replicate the style vector from the transformed image. (b) The model is additionally equipped with a cycle consistency loss which reconstructs the input image conditioned on the style of the transformed one. (c) A sketch of the working principles of IMPA. The model abstracts content from style and uses the latter to synthetically perturb an input control image to approximate the effect of a perturbation of interest.

**Supplementary Figure 2.**
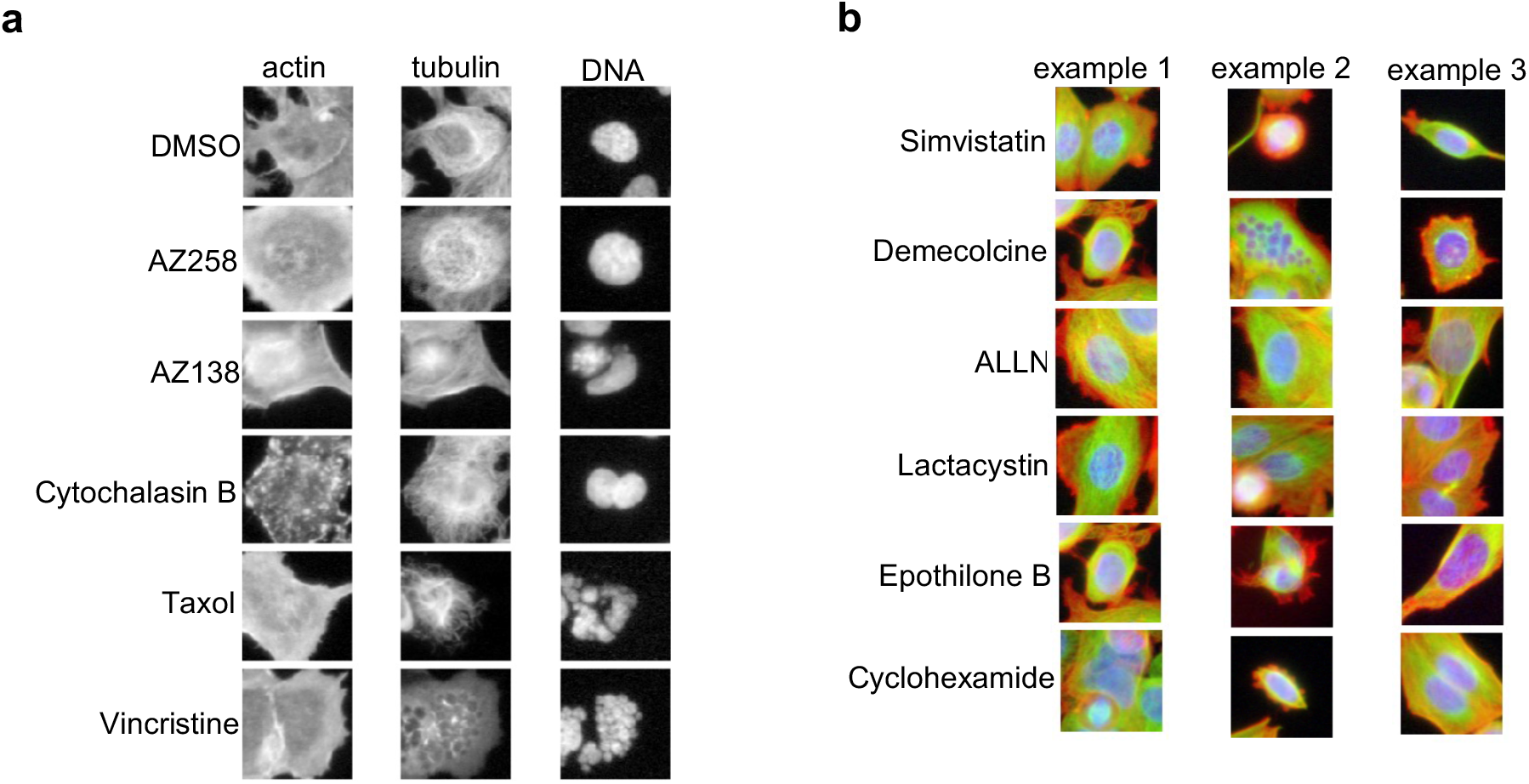
Examples of high-content images from BBBC021. **(a)** Examples of images of BBBC021 perturbed by 5 different drugs with distinct modes of action. Channels representing different substructures are displayed separately. An additional unperturbed cell instance (DMSO) is included as a reference. **(b)** Three examples of heterogenous perturbation effects by six drugs on cells from BBBC021.

**Supplementary Figure 3.**
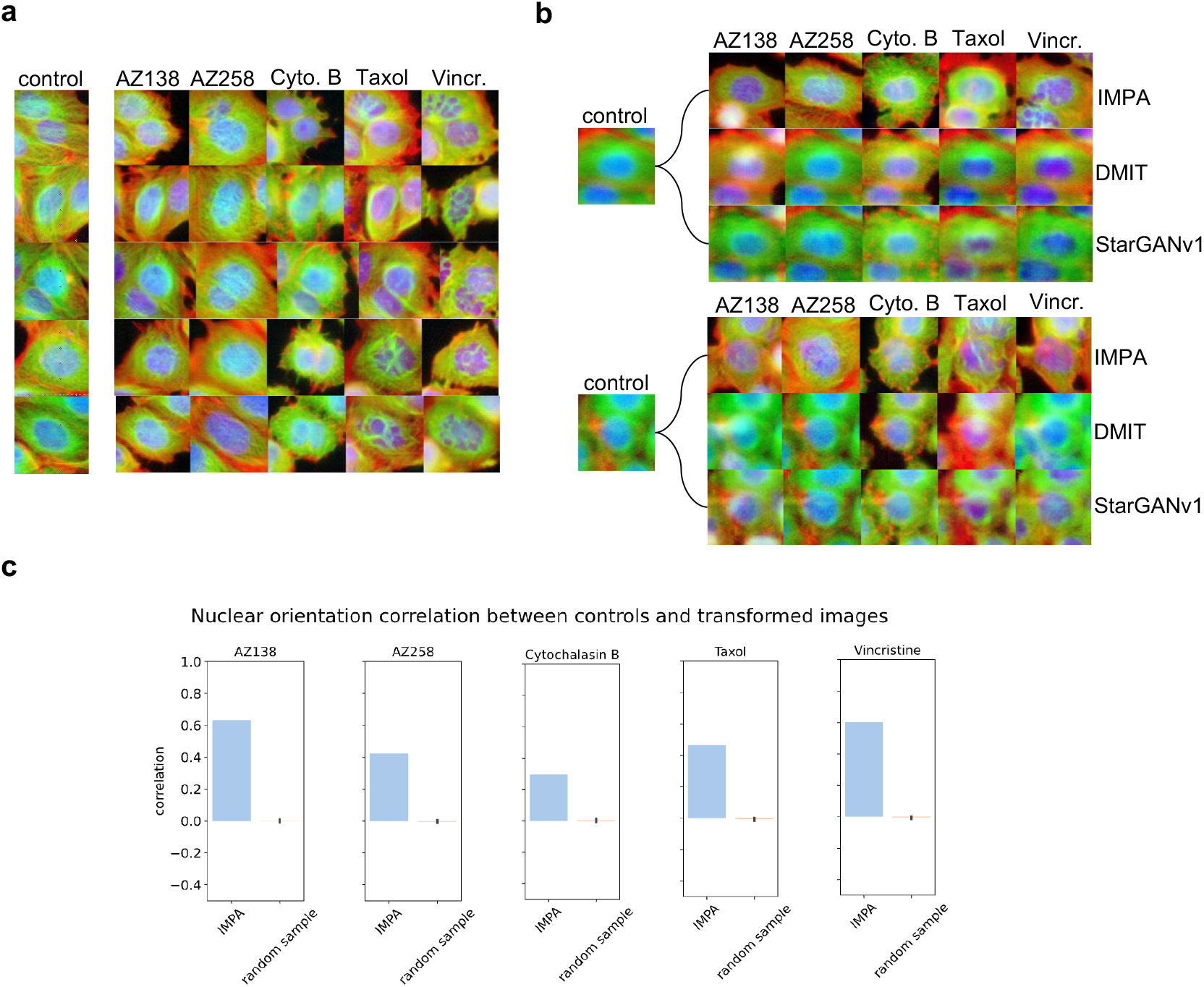
Visual exploration of transformations by IMPA. (a) Additional examples of cell transformations produced by IMPA on images from BBBC021 when trained with five drugs. (b) Comparison of predictions computed by IMPA and existing models on noisy inputs. (c) Correlation of nuclear orientations between the control images submitted to IMPA before and after transformation. A high correlation suggests that IMPA preserves cell orientation despite changing its style. Results are compared with the correlation between images of controls and random samples from the target drugs.

**Supplementary Figure 4.**
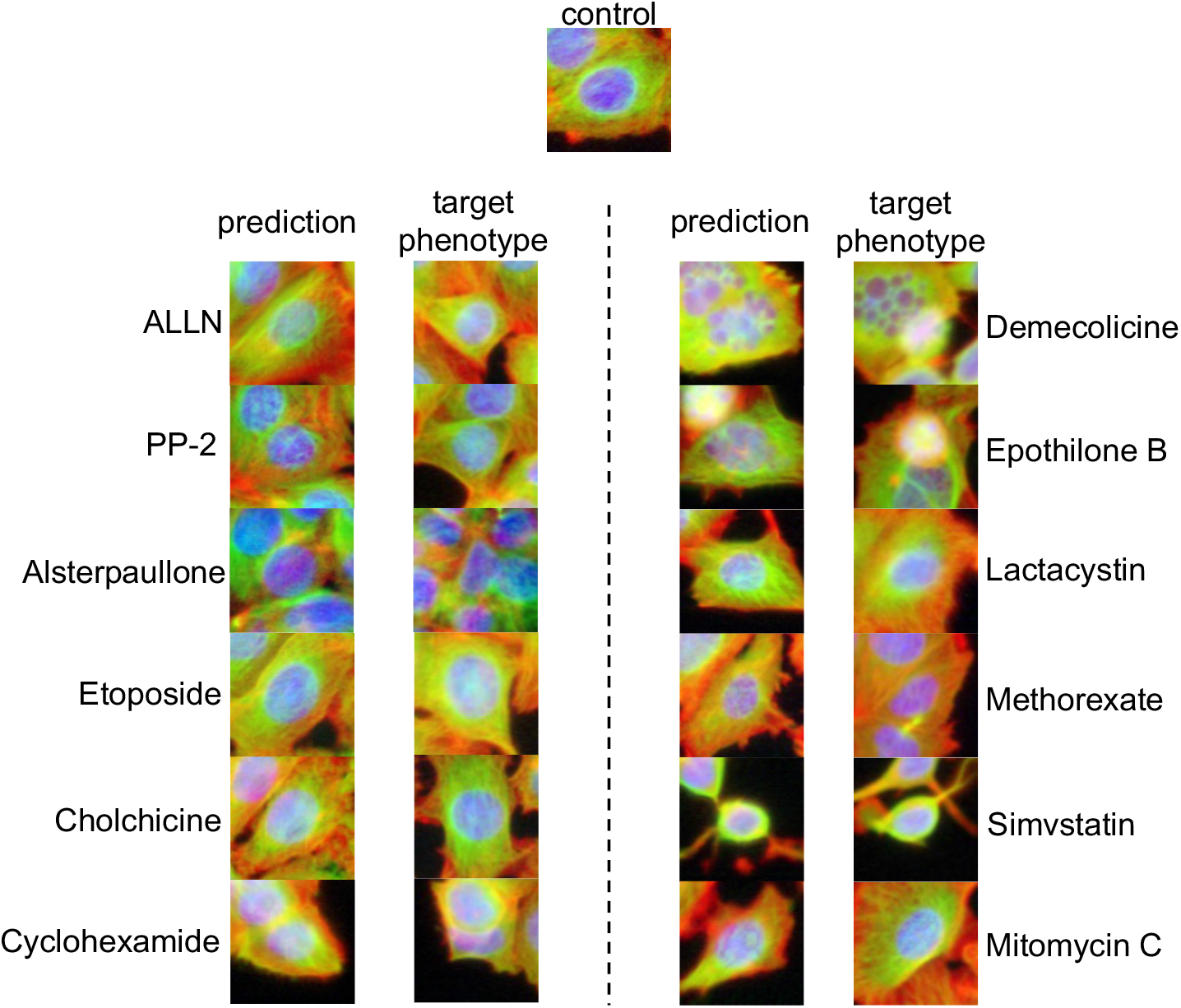
Extension of the predictions of IMPA to the whole BBBC021 dataset. Predictions by IMPA on 12 drugs after training on the whole BBBC021 dataset (34 drugs).

**Supplementary Figure 5.**
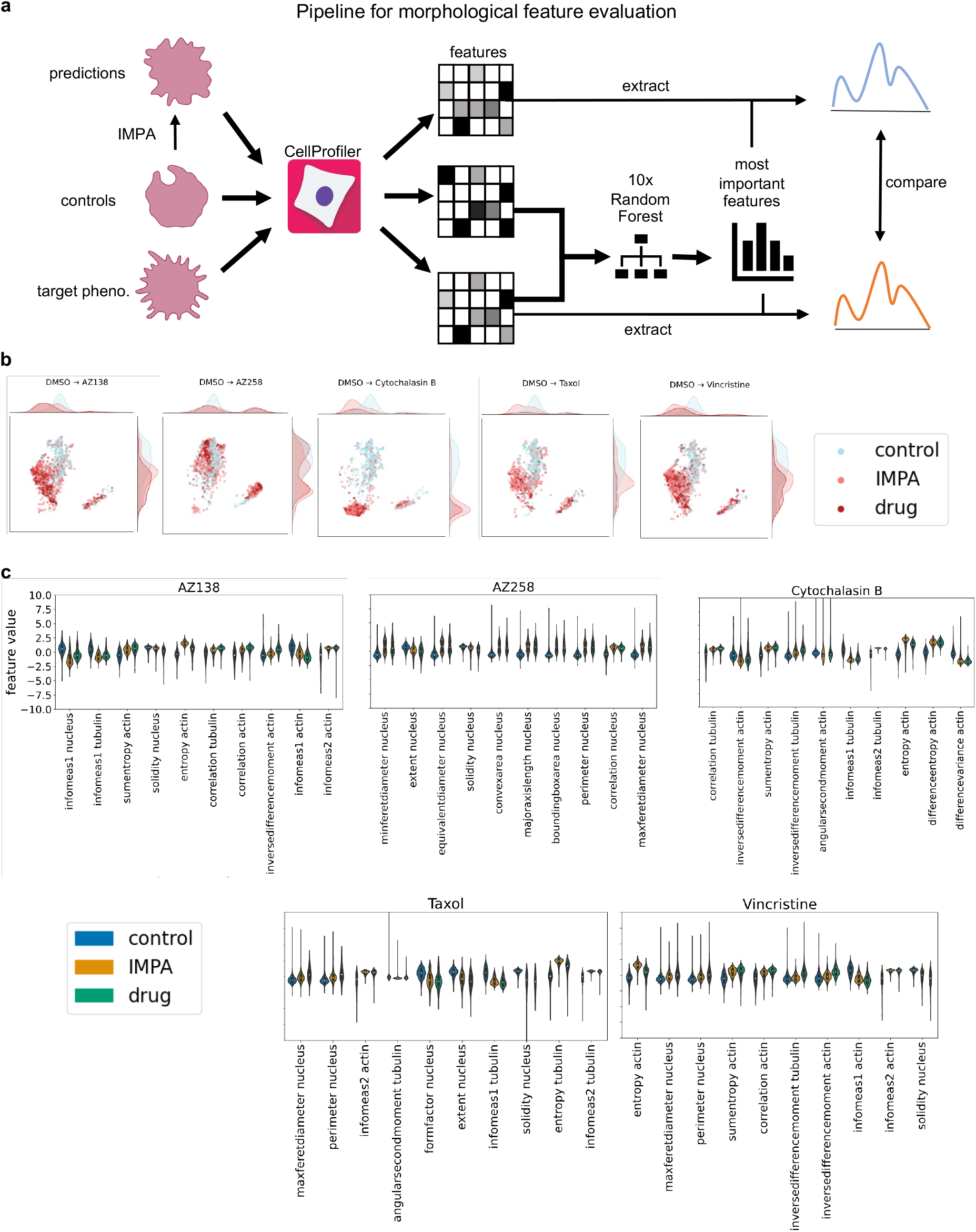
Morphological feature analysis with IMPA. **(a)** A schematic representation of the morphological feature analysis conducted to validate IMPA. Control images, synthetic predictions and real perturbation images are submitted to CellProfiler [51] to measure image features describing cell morphology. Feature importance is estimated via Random Forest comparing the real perturbed images with controls. Features extracted from real and predicted perturbations are compared to verify that the effect of IMPA on controls mimics measurable perturbation effects. **(b)** UMAP plots showing the distribution of morphological features before and after transformation with IMPA for five drugs included in the BBBC021 dataset. **(c)** Distribution of the ten most important discriminative features in controls, IMPA’s predictions and true perturbation images for five drugs in BBBC021.

**Supplementary Figure 6.**
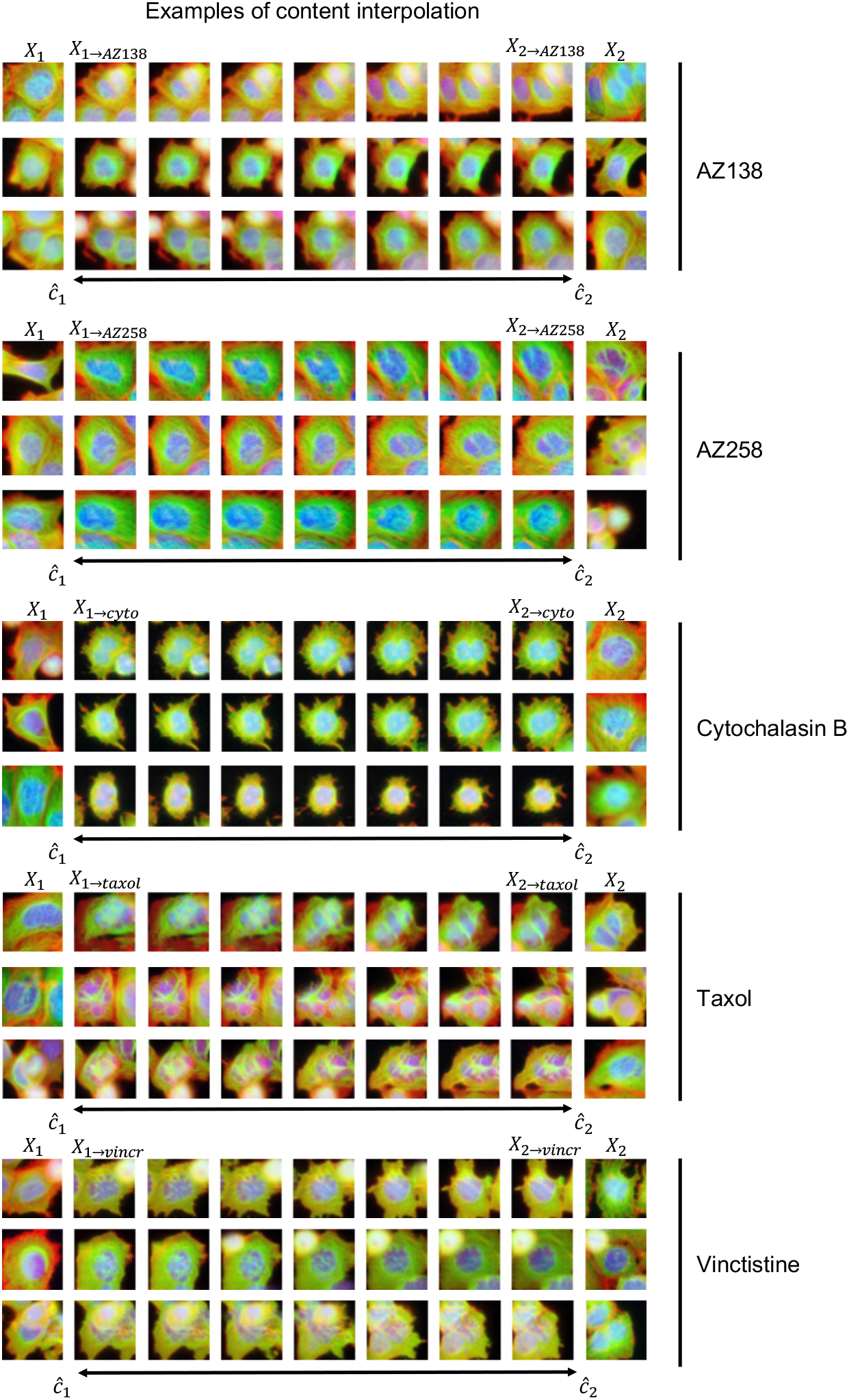
Content interpolation examples on BBBC021. Content interpolation of images of cells with constant styles. X_1_ and X_2_ are synthetically perturbed via IMPA by a chosen drug and their content encodings are interpolated to yield intermediate generated cells with fixed style.

**Supplementary Figure 7.**
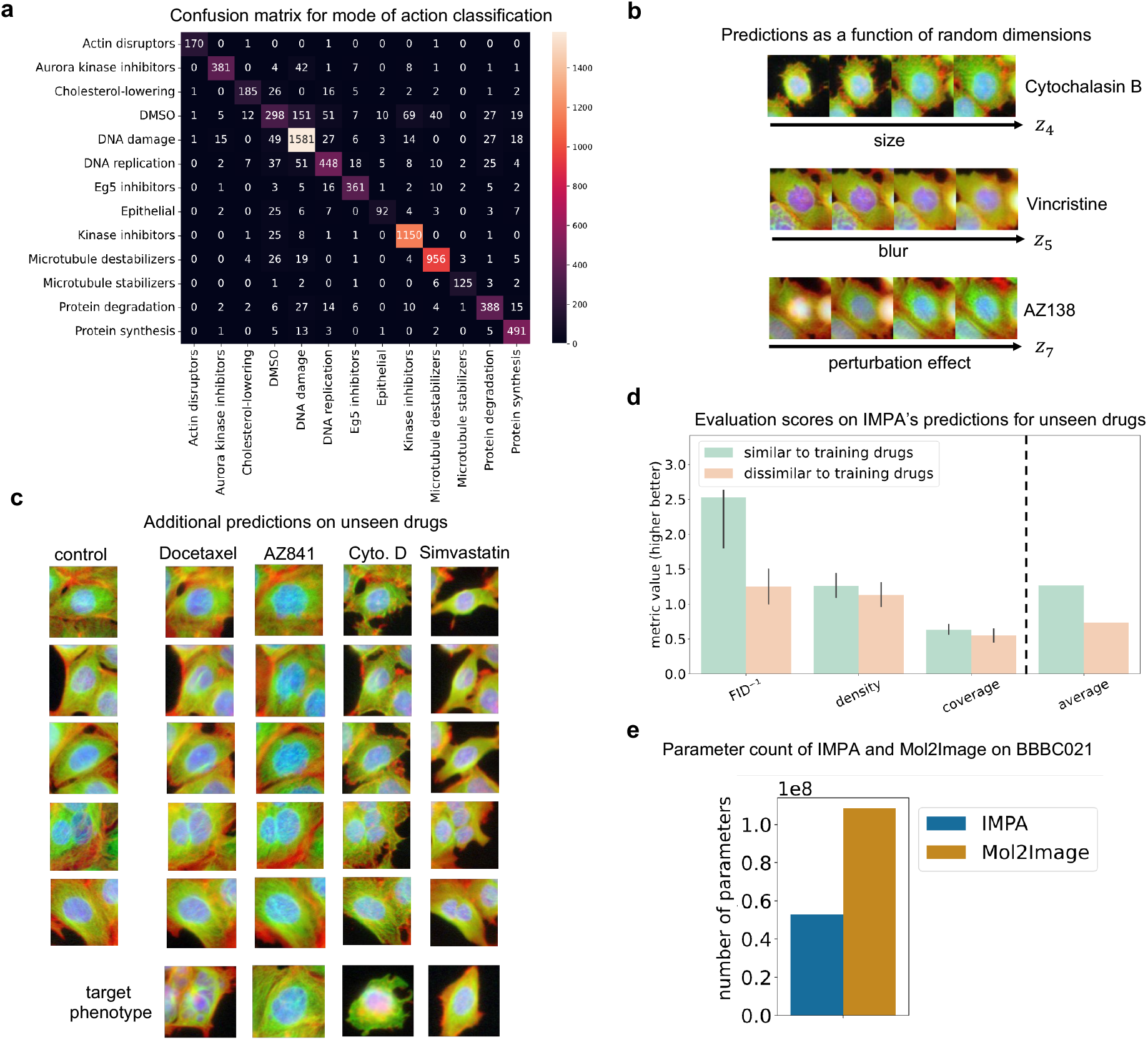
Additional experiments on BBBC021. **(a)** Confusion matrix for MoA prediction of the deep classifier trained to evaluate the output of IMPA. **(b)** Relation between generated morphological responses to drugs and single dimensions of the random Gaussian conditioning vector. **(c)** Additional examples of predictions on unseen drugs with a structure similar to training compounds. **(d)** Comparison of IMPA’s predictions on unseen drugs divided between similar and dissimilar to training compounds. For all measurements, the higher the value, the better the generated output approximates the expected phenotype. **(e)** The parameter count of IMPA and Mol2Image when run on the whole BBBC021 dataset.

**Supplementary Figure 8.**
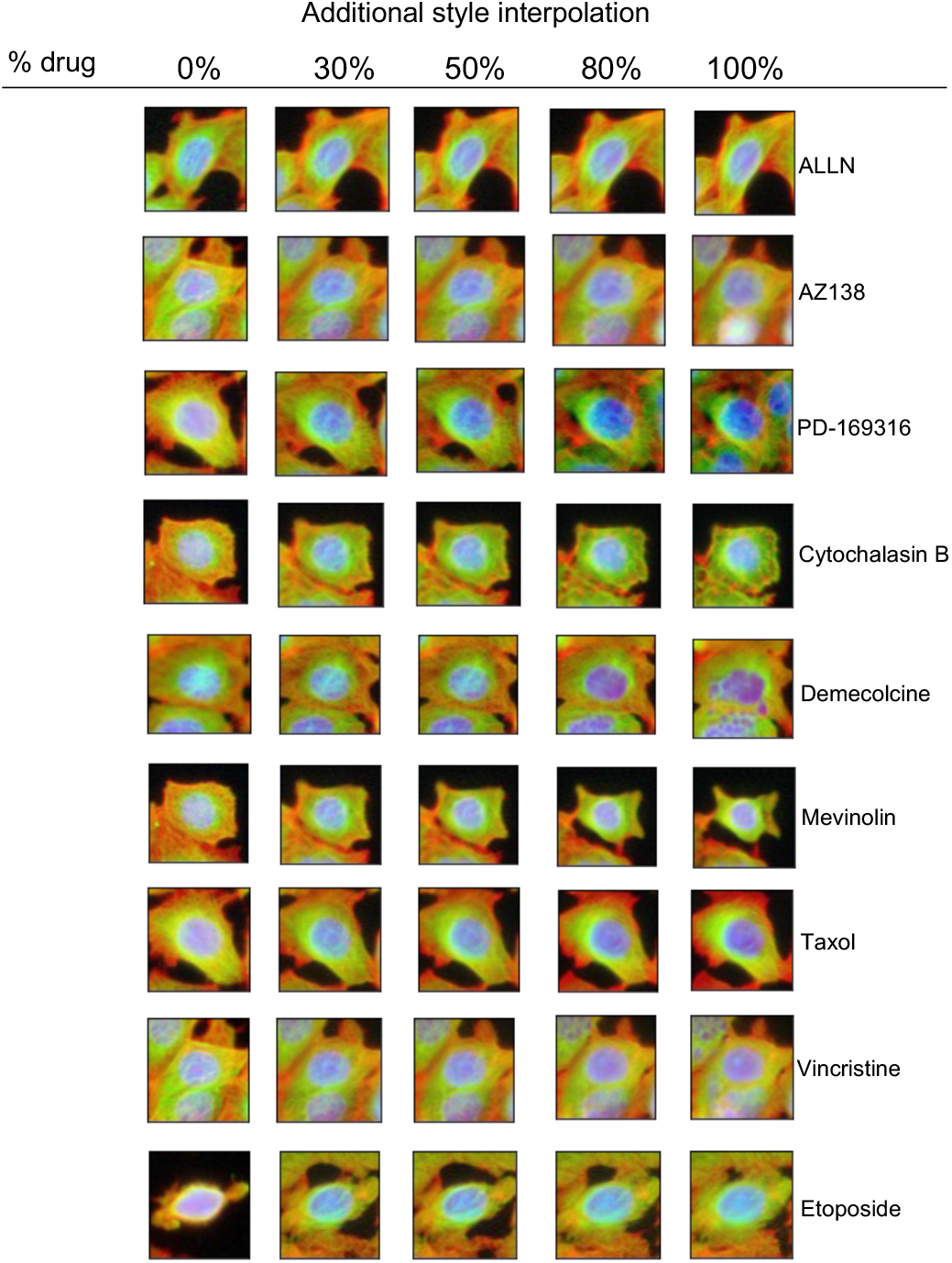
Style interpolation between control and a subset of drugs. Examples of style interpolation between controls and drugs. The content representation of images is first inferred for control cells and linearly interpolated with the style representation of target drugs at different intervals.

**Supplementary Figure 9.**
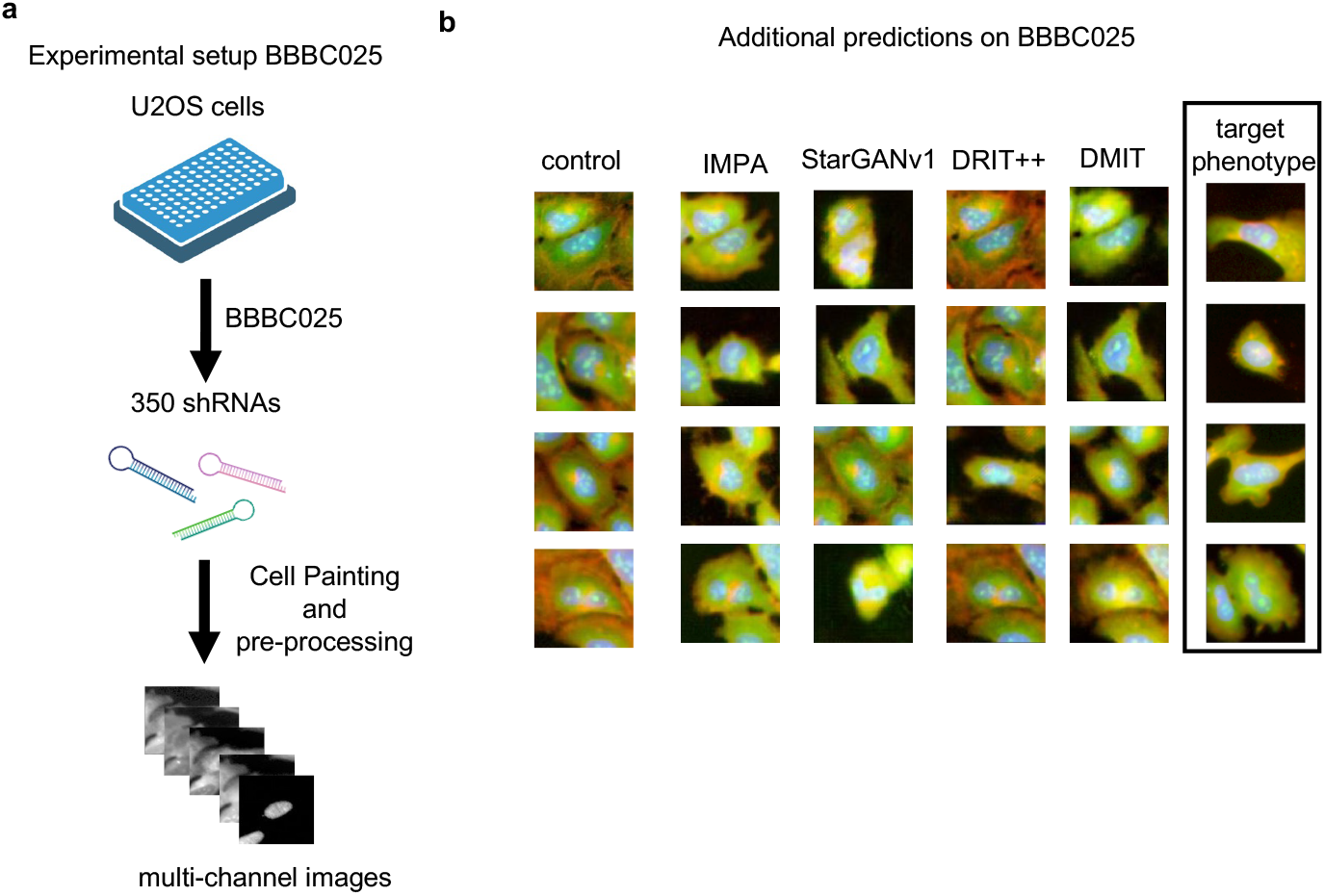
Prediction of active drug effects on BBBC025. (a) The experimental setup of data collection and processing on BBBC025. U2OS cells are plated with 350 different shRNA molecules targeting 41 distinct genes. The perturbed cells are subsequently imaged via the Cell Painting assay. (b) Comparison of active perturbation predictions between IMPA and existing methods on BBBC025.

**Supplementary Figure 10.**
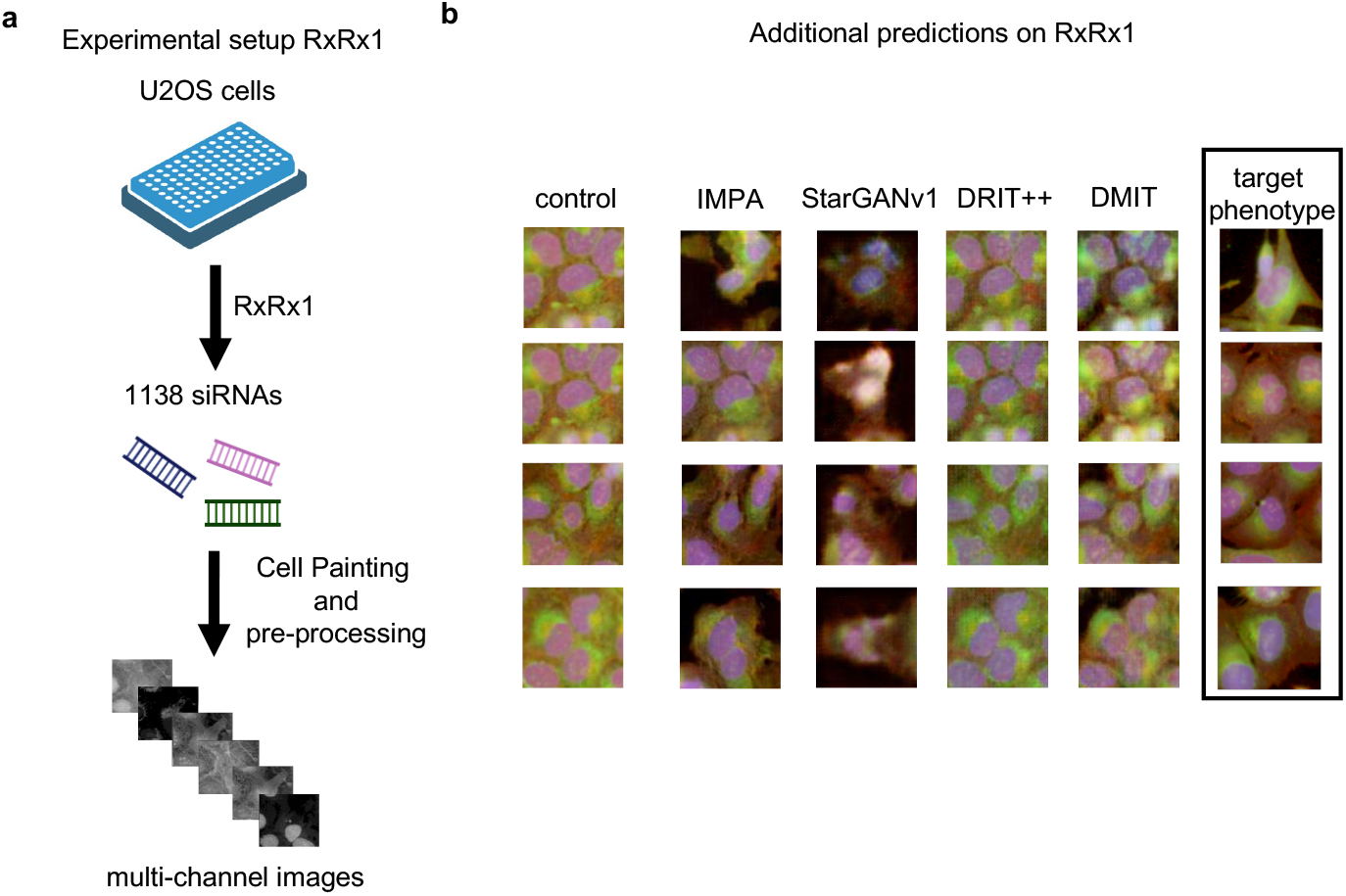
Prediction of active drug effects on RxRx1. (a) The experimental setup of data collection and processing on RxRx1. U2OS cells are plated with 1,138 different siRNA molecules. The perturbed cells are subsequently imaged via the Cell Painting assay. (b) Comparison of active perturbation predictions between IMPA and existing methods on RxRx1.

## References

[1] Bickle, M. The beautiful cell: high-content screening in drug discovery. Analytical and Bioanalytical Chemistry 398, 219–226 (2010).

[2] Boutros, M. et al. Microscopy-based high-content screening. Cell 163, 1314–1325 (2015).

[3] Lin, S. et al. Image-based high-content screening in drug discovery. Drug Discovery Today 25, 1348–1361 (2020).

[4] Moffat, J. G. et al. Opportunities and challenges in phenotypic drug discovery: an industry perspective. Nature Reviews Drug Discovery 16, 531–543 (2017).

[5] Zhou, Y. et al. High-throughput screening of a CRISPR/cas9 library for functional genomics in human cells. Nature 509, 487–491 (2014).

[6] Echeverri, C. J. et al. High-throughput RNAi screening in cultured cells: a user’s guide. Nature Reviews Genetics 7, 373–384 (2006).

[7] Scheeder, C. et al. Machine learning and image-based profiling in drug discovery. Current opinion in systems biology 10, 43–52 (2018).

[8] Bray, M.-A. et al. Cell painting, a high-content image-based assay for morphological profiling using multiplexed fluorescent dyes. Nature Protocols 11, 1757–1774 (2016).

[9] Chandrasekaran, S. N. et al. Image-based profiling for drug discovery: due for a machine-learning upgrade? Nature Reviews Drug Discovery 20, 145–159 (2021).

[10] Schimunek, J., et al. Context-enriched molecule representations improve few-shot drug discovery (2023).

[11] Polishchuk, P. G. et al. Estimation of the size of drug-like chemical space based on GDB-17 data. Journal of Computer-Aided Molecular Design 27, 675–679 (2013).

[12] Moffat, J. G. et al. Phenotypic screening in cancer drug discovery—past, present and future. Nature reviews Drug discovery 13, 588–602 (2014).

[13] Ljosa, V. et al. Comparison of methods for image-based profiling of cellular morphological responses to small-molecule treatment. Journal of biomolecular screening 18, 1321–1329 (2013).

[14] Ando, D. M., et al. Improving phenotypic measurements in high-content imaging screens. Cold Spring Harbor Laboratory (2017).

[15] Pawlowski, N., et al. Automating morphological profiling with generic deep convolutional networks. *bioRxiv* (2016).

[16] Perakis, A., et al. Contrastive learning of single-cell phenotypic representations for treatment classification. *arXiv* (2021).

[17] Godinez, W. J. et al. A multi-scale convolutional neural network for phenotyping high-content cellular images. Bioinformatics 33, 2010–2019 (2017).

[18] Kensert, A. et al. Transfer learning with deep convolutional neural networks for classifying cellular morphological changes. SLAS Discovery 24, 466–475 (2019).

[19] Nyffeler, J. et al. Bioactivity screening of environmental chemicals using imaging-based high-throughput phenotypic profiling. Toxicology and Applied Pharmacology 389, 114876 (2020).

[20] Hofmarcher, M. et al. Accurate prediction of biological assays with high-throughput microscopy images and convolutional networks. Journal of Chemical Information and Modeling 59, 1163–1171 (2019).

[21] Lafarge, M. W. et al. Capturing single-cell phenotypic variation via unsupervised representation learning. In Cardoso, M. J. et al. (eds.) Proceedings of The 2nd International Conference on Medical Imaging with Deep Learning, vol. 102 of Proceedings of Machine Learning Research, 315–325 (PMLR, 2019).

[22] Chow, Y. L. et al. Predicting drug polypharmacology from cell morphology readouts using variational autoencoder latent space arithmetic. PLoS computational biology 18, e1009888 (2022).

[23] Lee, H., et al. MorphNet predicts cell morphology from single-cell gene expression. *bioRxiv* (2022).

[24] Klambauer, G., et al. CLOOME: contrastive learning unlocks bioimaging databases for queries with chemical structures (2022).

[25] Pernice, W. M. et al. Out of distribution generalization via interventional style transfer in single-cell microscopy. In Proceedings of the IEEE/CVF Conference on Computer Vision and Pattern Recognition (CVPR) Workshops, 4325–4334 (2023).

[26] Yang, K. et al. Mol2image: Improved conditional flow models for molecule to image synthesis. In Proceedings of the IEEE/CVF Conference on Computer Vision and Pattern Recognition, 6688–6698 (2021).

[27] Gatys, L. A. et al. Image style transfer using convolutional neural networks. In Proceedings of the IEEE Conference on Computer Vision and Pattern Recognition (CVPR) (2016).

[28] Jing, Y. et al. Neural style transfer: A review. IEEE Transactions on Visualization and Computer Graphics 26, 3365–3385 (2020).

[29] Li, Y., et al. Demystifying neural style transfer. *arXiv* (2017).

[30] Zhang, Y. et al. Separating style and content for generalized style transfer. In Proceedings of the IEEE Conference on Computer Vision and Pattern Recognition (CVPR) (2018).

[31] Pang, Y. et al. Image-to-image translation: Methods and applications. IEEE Transactions on Multimedia 24, 3859–3881 (2021).

[32] Isola, P. et al. Image-to-image translation with conditional adversarial networks. arXiv (2016).

[33] Huang, X., et al. Arbitrary style transfer in real-time with adaptive instance normalization. *arXiv* (2017).

[34] Jing, Y. et al. Neural style transfer: A review. IEEE Transactions on Visualization and Computer Graphics 26, 3365–3385 (2020).

[35] Emami, H. et al. SPA-GAN: Spatial attention GAN for image-to-image translation. IEEE Transactions on Multimedia 23, 391–401 (2021).

[36] Sauer, A., et al. StyleGAN-T: Unlocking the power of GANs for fast large-scale text-to-image synthesis. *arXiv* (2023).

[37] Mirza, M., et al. Conditional generative adversarial nets. *arXiv* (2014).

[38] Choi, Y. et al. Stargan v2: Diverse image synthesis for multiple domains. arXiv (2019).

[39] Landrum, G. Rdkit: Open-source cheminformatics software (2016).

[40] Du, J. et al. Gene2vec: distributed representation of genes based on co-expression. BMC Genomics 20, 82 (2019).

[41] Hetzel, L., et al. Predicting single-cell perturbation responses for unseen drugs. In ICLR2022 Machine Learning for Drug Discovery (2022).

[42] Lotfollahi, M. et al. Biologically informed deep learning to query gene programs in single-cell atlases. Nature Cell Biology 25, 337–350 (2023).

[43] Yu, H., et al. Perturbnet predicts single-cell responses to unseen chemical and genetic perturbations. *bioRxiv* (2022).

[44] Lotfollahi, M. et al. Predicting cellular responses to complex perturbations in high-throughput screens. Molecular Systems Biology e11517 (2023).

[45] Goodfellow, I. J. et al. Generative adversarial networks. arXiv (2014).

[46] Zhu, J.-Y., et al. Unpaired image-to-image translation using cycle-consistent adversarial networks. *arXiv* (2017).

[47] Ljosa, V. et al. Comparison of methods for image-based profiling of cellular morphological responses to small-molecule treatment. SLAS Discovery 18, 1321–1329 (2013). Special Issue: Phenotypic Drug Discovery (Part 1 of 2).

[48] Lo, Y.-C. et al. Machine learning in chemoinformatics and drug discovery. Drug Discovery Today 23, 1538–1546 (2018).

[49] Kulms, D. et al. Apoptosis induced by disruption of the actin cytoskeleton is mediated via activation of CD95 (fas/APO-1). Cell Death &amp Differentiation 9, 598–608 (2002).

[50] Huang, Y. et al. Regulation of ivinca/i alkaloid-induced apoptosis by NF-b/ib pathway in human tumor cells. Molecular Cancer Therapeutics 3, 271–277 (2004).

[51] McQuin, C. et al. Cellprofiler 3.0: Next-generation image processing for biology. PLOS Biology 16, 1–17 (2018).

[52] Ho, T. K. Random decision forests. In Proceedings of 3rd international conference on document analysis and recognition, vol. 1, 278–282 (IEEE, 1995).

[53] Choi, Y. et al. Stargan: Unified generative adversarial networks for multi-domain image-to-image translation. In Proceedings of the IEEE Conference on Computer Vision and Pattern Recognition (CVPR) (2018).

[54] Lee, H.-Y., et al. Drit++: Diverse image-to-image translation via disentangled representations. *arXiv* (2019).

[55] Yu, X., et al. Multi-mapping image-to-image translation via learning disentanglement. In NeurIPS (2019).

[56] Pang, Y. et al. Image-to-image translation: Methods and applications. IEEE Transactions on Multimedia 24, 3859–3881 (2021).

[57] Heusel, M. et al. Gans trained by a two time-scale update rule converge to a local nash equilibrium. In Proceedings of the 31st International Conference on Neural Information Processing Systems, NIPS’17, 6629–6640 (Curran Associates Inc., Red Hook, NY, USA, 2017).

[58] Naeem, M. F. et al. Reliable fidelity and diversity metrics for generative models. In III, H. D. & Singh, A. (eds.) Proceedings of the 37th International Conference on Machine Learning, vol. 119 of Proceedings of Machine Learning Research, 7176–7185 (PMLR, 2020).

[59] Maragakis, P. et al. A deep-learning view of chemical space designed to facilitate drug discovery. Journal of Chemical Information and Modeling 60, 4487–4496 (2020).

[60] Li, X. et al. Chemical space exploration based on recurrent neural networks: applications in discovering kinase inhibitors. Journal of Cheminformatics 12 (2020).

[61] Blanco-Gonzalez, A., et al. The role of ai in drug discovery: Challenges, opportunities, and strategies. *arXiv* (2022).

[62] Vukicevic, S. Current challenges and hurdles in new drug development. Clinical Therapeutics 38, e3 (2016).

[63] Szegedy, C. et al. Rethinking the inception architecture for computer vision. In Proceedings of the IEEE Conference on Computer Vision and Pattern Recognition (CVPR) (2016).

[64] Chandrasekaran, S. N. et al. JUMP cell painting dataset: morphological impact of 136, 000 chemical and genetic perturbations (2023).

[65] Gilmer, J., et al. Neural message passing for quantum chemistry. *arXiv* (2017).

[66] Gómez-Bombarelli, R. et al. Automatic chemical design using a data-driven continuous representation of molecules. ACS Central Science 4, 268–276 (2018).

[67] Jin, W. et al. Junction tree variational autoencoder for molecular graph generation. In Dy, J. & Krause, A. (eds.) Proceedings of the 35th International Conference on Machine Learning, vol. 80 of Proceedings of Machine Learning Research, 2323–2332 (PMLR, 2018).

[68] Kearnes, S. et al. Molecular graph convolutions: moving beyond fingerprints. Journal of Computer-Aided Molecular Design 30, 595–608 (2016).

[69] Rong, Y., et al. Self-supervised graph transformer on large-scale molecular data. In Larochelle, H., Ranzato, M., Hadsell, R., Balcan, M. & Lin, H. (eds.) Advances in Neural Information Processing Systems, vol. 33, 12559–12571 (Curran Associates, Inc., 2020).

[70] Ho, J., et al. Denoising diffusion probabilistic models (2020).

[71] Mao, Q. et al. Mode seeking generative adversarial networks for diverse image synthesis. In Proceedings of the IEEE/CVF Conference on Computer Vision and Pattern Recognition (CVPR) (2019).

[72] Ulyanov, D., et al. Instance normalization: The missing ingredient for fast stylization. *arXiv* (2016).

[73] Ioffe, S. et al. Batch normalization: Accelerating deep network training by reducing internal covariate shift. In Bach, F. & Blei, D. (eds.) Proceedings of the 32nd International Conference on Machine Learning, vol. 37 of Proceedings of Machine Learning Research, 448–456 (PMLR, Lille, France, 2015).

[74] He, K. et al. Deep residual learning for image recognition. In Proceedings of the IEEE Conference on Computer Vision and Pattern Recognition (CVPR) (2016).

[75] He, K. et al. Delving deep into rectifiers: Surpassing human-level performance on imagenet classification. In Proceedings of the IEEE International Conference on Computer Vision (ICCV) (2015).

[76] Kingma, D. P., et al. Glow: Generative flow with invertible 1×1 convolutions. In Bengio, S., et al. (eds.) Advances in Neural Information Processing Systems, vol. 31 (Curran Associates, Inc., 2018).

[77] Deng, J. et al. Imagenet: A large-scale hierarchical image database. In 2009 IEEE Conference on Computer Vision and Pattern Recognition, 248–255 (2009).

[78] Otsu, N. A threshold selection method from gray-level histograms. *IEEE Transactions on Systems*, Man, and Cybernetics 9, 62–66 (1979).

[79] Roerdink, J. B. T. M. et al. The watershed transform: Definitions, algorithms and parallelization strategies. Fundam. Inf. 41, 187–228 (2000).

[80] Caie, P. D. et al. High-Content Phenotypic Profiling of Drug Response Signatures across Distinct Cancer Cells. Molecular Cancer Therapeutics 9, 1913–1926 (2010).

[81] Singh, S. et al. Morphological profiles of RNAi-induced gene knockdown are highly reproducible but dominated by seed effects. PLOS ONE 10, e0131370 (2015).

[82] Bray, M.-A. et al. Cell painting, a high-content image-based assay for morphological profiling using multiplexed fluorescent dyes. Nature Protocols 11, 1757–1774 (2016).

[83] Taylor, J., et al. Rxrx1: an image set for cellilar morphological variation across many experimental batches. ICLR AI for social good workshop (2019).

[84] Singh, S., et al. Pipeline for illumination correction of images for high-throughput microscopy. Journal of Microscopy 256, 231–236 (2014).

[85] Crété-Roffet, F., et al. The Blur Effect: Perception and Estimation with a New No-Reference Perceptual Blur Metric. In SPIE Electronic Imaging Symposium Conf Human Vision and Electronic Imaging, vol. XII, EI 6492–16 (San Jose, United States, 2007).

[86] Mikolov, T., et al. Efficient estimation of word representations in vector space (2013).

